# Ndc80 complex, a conserved coupler for kinetochore-microtubule motility, is a sliding molecular clutch

**DOI:** 10.1101/2025.03.13.643154

**Authors:** Vladimir M. Demidov, Ivan V. Gonchar, Suvranta K. Tripathy, Fazly I. Ataullakhanov, Ekaterina L. Grishchuk

**Affiliations:** Department of Physiology, University of Pennsylvania; PA, United States

**Author notes:** equal contribution (Vladimir M. Demidov, Ivan V. Gonchar). Department of Natural Sciences, University of Michigan-Dearborn; Dearborn, MI, USA.

**Keywords:** chromosome segregation, force spectroscopy, kinetochore slip-clutch, microtubule cytoskeleton, molecular friction, optical trapping, Brownian model

## Abstract

Chromosome motion at spindle microtubule plus-ends relies on dynamic molecular bonds between kinetochores and proximal microtubule walls. Under opposing forces, kinetochores move bi-directionally along these walls while remaining near the ends, yet how continuous wall-sliding occurs without end-detachment remains unclear. Using ultrafast force-clamp spectroscopy, we show that single Ndc80 complexes, the primary microtubule-binding kinetochore component, exhibit processive, bi-directional sliding. Plus-end-directed forces induce a mobile catch-bond in Ndc80, increasing frictional resistance and restricting sliding toward the tip. Conversely, forces pulling Ndc80 away from the plus-end trigger mobile slip-bond behavior, facilitating sliding. This dual behavior arises from force-dependent modulation of the Nuf2 calponin-homology domain’s microtubule binding, identifying this subunit as a friction regulator. We propose that Ndc80c’s ability to modulate sliding friction provides the mechanistic basis for the kinetochore’s end coupling, enabling its slip-clutch behavior.

**One Sentence Summary:** Direction-dependent mobile catch- and slip-bond behavior of the microtubule-binding Ndc80 protein

## Introduction

### Slip-Clutch Behavior Modulates Friction at the Kinetochore-Microtubule Interface for Accurate Chromosome Segregation

During cell division, kinetochores establish intricate attachments near the plus-ends of spindle microtubules, enabling chromosome segregation through coordinated motility *(1-3)*. In metaphase of human cells, depolymerization of kinetochore-bound microtubules alternates at sister kinetochores, generating pulling forces that drive chromosome oscillations between spindle poles. Depolymerization at the “leading” kinetochore pulls the attached chromosome toward the pole, while the “trailing” kinetochore passively follows (Fig. 1A) *(1, 2, 4-7)*. This motion is enabled by the phenomenon referred to as end-coupling, during which kinetochore slides along end-proximal microtubule walls via a differential frictional interface, functioning as a slip-clutch *(8-11)*. The trailing kinetochore, dragged along polymerizing microtubules toward their plus-ends, exhibits clutch-like behavior that protects it from end detachment, while reduced friction at the leading kinetochore facilitates its pole-directed sliding, driven by the depolymerization motor *(12, 13)*. Similar frictional regulation may also assist poleward kinetochore motion in anaphase while resisting forces that could pull the kinetochore off the plus-end. The biophysical and molecular basis underlying such regulation remain unresolved.

**Figure 1.**
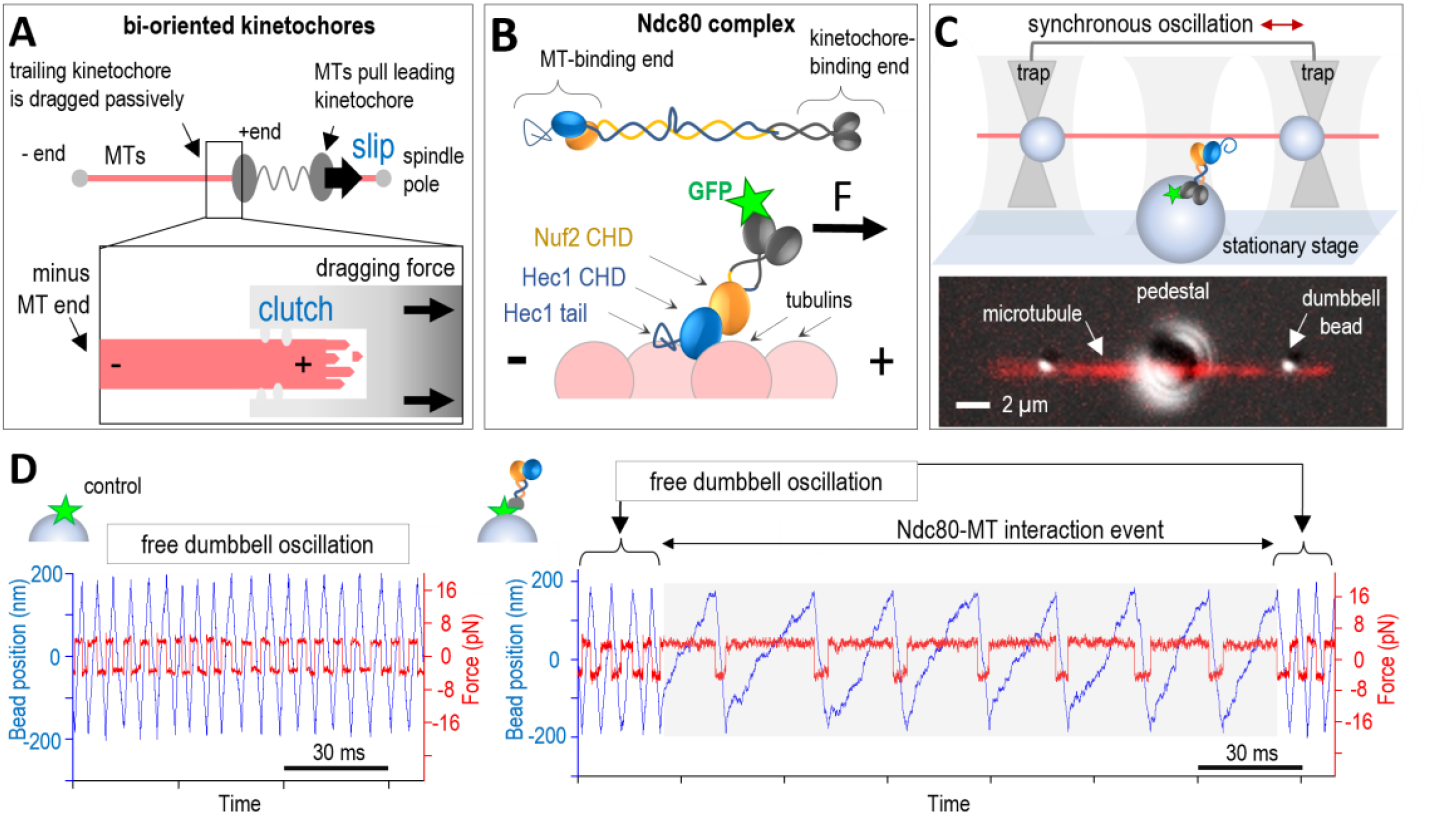
Bi-directional processive sliding of single Ndc80c molecules revealed by the UFFC assay. (**A**) Kinetochores slide along the microtubule wall in alternating directions, transitioning between the highly mobile “slip” state and the “clutch” state, which generates higher molecular friction to protect the trailing kinetochore from detachment at the microtubule plus-end. (**B**) Ndc80c binds microtubule-wall with the “toe” of the Hec1 calponin-homology domain (CHD) with a pronounced tilt toward the microtubule plus-end. (**C**) Schematic of the three-bead assay setup and a representative image showing merged fluorescence and DIC (Differential Interference Contrast) channels. (**D**) Representative signals showing changes in the position of a dumbbell bead (blue) and the clamped force (red) during experiments with GFP- or Ndc80c-coated pedestals. Grey areas correspond to continuous Ndc80c-microtubule interaction.

Microtubule-end coupling is mediated by the Ndc80 protein complex (Ndc80c), a key component of the kinetochore-microtubule interface and a crucial player in chromosome segregation *(14-16)*. At the kinetochore, this microtubule wall-binding protein forms a molecular “lawn”, where individual Ndc80c molecules dynamically associate and dissociate from microtubules *(17, 18)*. A widely accepted model for the mechanism of end-coupling suggests that Ndc80c translocates along microtubules via biased diffusion *(19-21)*. However, a purely diffusive mechanism is inconsistent with the observed slip-clutch kinetochore behavior, as it predicts a direction-independent frictional interface. Thus, additional molecular mechanisms beyond biased diffusion must contribute to kinetochore-microtubule coupling. Whether individual Ndc80c molecules exhibit direction-dependent slip-clutch dynamics remains an open question, central to understanding the molecular basis of kinetochore-microtubule coupling.

Ndc80c binds microtubules through the N-terminus of the Hec1 subunit, an unstructured extension that lacks a defined tubulin interaction footprint *(22-24)*. Additionally, Ndc80’s microtubule-binding end contains two closely associated calponin-homology domains (CHDs) from the Hec1 and Nuf2 subunits *(25, 26)*. Cryo-EM structures of densely packed Ndc80 complexes along microtubules reveal that the “toe” of the Hec1 CHD forms the sole site-specific binding interface with polymerized tubulins, while the tightly coupled Nuf2 CHD hovers near the tubulin surface without direct contact (Fig. 1B) *(27*). Interestingly, mutational analysis of Hec1 and Nuf2 CHDs in mitotic cells has shown that both contribute to microtubule attachment, with Nuf2 CHD playing an important role in generating tension across sister kinetochores *(28)*. The Nuf2 CHD may facilitate microtubule coupling indirectly by serving as a hub for interactions with mitotic regulators *(29-31)*. However, charge-altering mutations in this domain reduce Ndc80c’s affinity for microtubule walls in vitro *(25)*, raising questions about the physiological significance of its microtubule binding. Here, we investigated Ndc80c-microtubule interactions under force, uncovering a critical role of the Nuf2 CHD in enabling slip-clutch behavior of this essential kinetochore component.

## Results

### Ultrafast force-clamp spectroscopy uncovers force-guided sliding of single Ndc80c molecules

In the absence of external force, single Ndc80c molecules transiently bind to microtubules for fraction of a second *(32, 33)*. Investigating the load dependence of such short-lived interactions in non-motor proteins requires advanced single-molecule techniques, such as ultrafast force-clamp (UFFC) spectroscopy *(34-36)*. The UFFC assay employs a “three-bead” geometry, where a taxol-stabilized microtubule is suspended between two beads held in optical traps (Fig. 1C). This microtubule dumbbell is oscillated along its axis by synchronously driving the traps with a 30 MHz acousto-optic deflector *(35)*. Viscous drag on the dumbbell enables continuous force-clamping, with the applied force reversing direction every 0.4 µm. When the oscillating dumbbell is positioned near a coverslip-immobilized “pedestal” bead coated with Ndc80c molecules, binding to the moving microtubule subjects the Ndc80c to directional forces ultrafast—within just 50 µs of attachment. To preserve Ndc80c’s natural orientation, it was immobilized via a GFP tag at its kinetochore-binding end *(25)*. The “Bonsai” variant of human Ndc80 was used, retaining the complete microtubule-binding interface but featuring a significantly shortened rod (< 20 nm) to reduce linkage compliance and length-dependent effects *(25)*

During UFFC assay, Ndc80c binding events were detected through changes in dumbbell bead positions, monitored by two detection beams and quadrant photodetectors. When dumbbells oscillated near sparsely coated Ndc80c pedestals, 19% exhibited deviations from the regular oscillation pattern, consistent with single-molecule binding events (4 pN clamp force, n=314, Table S1). In contrast, GFP-coated pedestals did not disrupt oscillations, and microtubule dumbbells maintained their characteristic “free” velocity, confirming that GFP alone does not interact with the microtubule (Fig. 1D). If individual Ndc80c-microtubule bonds lacked sliding capacity, after Ndc80 binding the force clamp would cause dumbbell motion to cease, maintaining force stably until bond dissociation (Fig. S2). In molecular ensembles, such transient interactions would produce overall frictional resistance despite the absence of sliding capability at the single-molecule level. However, no such pausing events were observed for Ndc80c. Instead, upon Ndc80c binding, the dumbbell motion continued, albeit at reduced velocities, indicating sustained microtubule contact (Fig. 1D, Fig. S3). During each 400-nm unidirectional sweep, Ndc80c translocated across arrays of up to 100 tubulin monomers. Given the 4-nm spacing of Ndc80c binding sites, this behavior unequivocally demonstrates processive, force-induced sliding by individual Ndc80c molecules.

### Polarity-Dependent Sliding is an Intrinsic Property of Ndc80c’s Microtubule-Binding Domains

During single microtubule-binding events, Ndc80c slid repeatedly in opposite directions under alternating pulling forces (Fig. 2A; Figs. S3-S7; Supplemental Materials and Methods). Velocity distributions for each microtubule-pedestal pair revealed a pronounced asymmetry: significantly slower velocity segments were confined to one pulling direction, while the opposite direction exhibited only minor retardation relative to the free dumbbell velocity (Fig. 2A). Ndc80c sliding at different velocities under the same pulling force suggests that it resists movement more strongly in one direction, with slower motion indicating greater resistive friction. Testing a single dumbbell against multiple Ndc80c-coated pedestals consistently reproduced the same asymmetry, indicating that the effect is not due to specific properties of the pedestal-immobilized Ndc80c but may instead arise from microtubule polarity *(37)*. To confirm this property at the single-molecule level, we introduced additional pedestals coated with either kinesin or dynein motors, proteins with well-established directional preferences on microtubule walls. The same dumbbell was first oscillated near an Ndc80c-coated pedestal and then suspended near a motor-coated pedestal in ATP-containing buffer to record motor-driven displacements (Fig. 2B). Kinesin consistently pulled the microtubule in the same direction as slow Ndc80c sliding, whereas dynein pulled in the direction of fast sliding (n = 5 and 9, respectively). These findings demonstrate that single Ndc80c molecules generate greater molecular friction under plus-end-directed dragging forces, aligning with predictions for kinetochore motility *(9, 10, 38)*.

**Figure 2.**
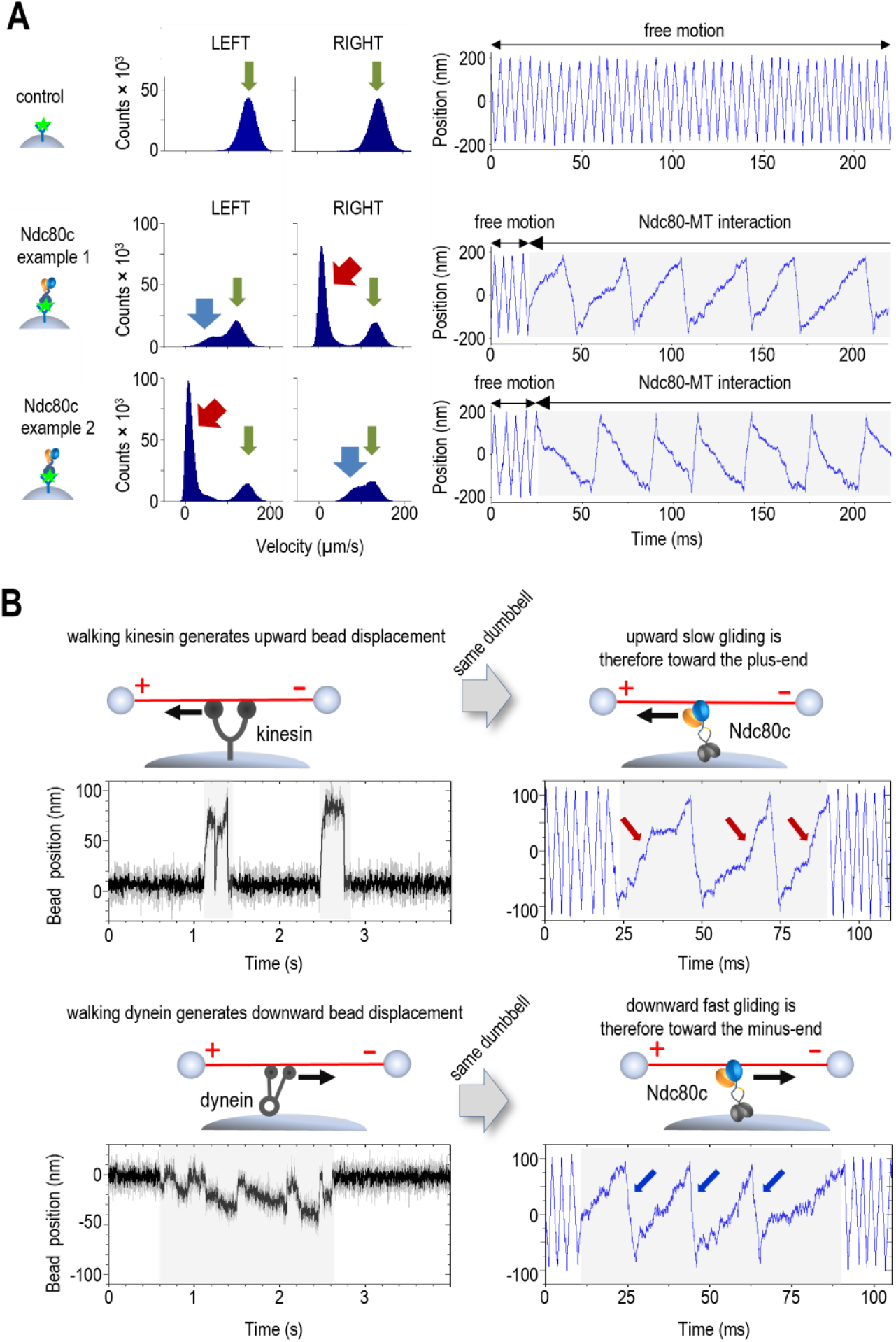
Sliding friction of single Ndc80c molecules is strongly microtubule-polarity dependent. (**A**) Histograms showing distributions of instantaneous velocities in two oscillation directions along with typical coordinate recordings of the dumbbell beads for 30 s at 4 pN clamp force. Rightward dumbbell oscillation corresponds to an upward change in bead coordinate on the graphs, while leftward dumbbell motion results in a decreasing bead coordinate. The first row depicts a control experiment with GFP-coated pedestals, showing “free” velocity peaks in two directions (green arrows). The following rows show results from two different dumbbells oscillated near Ndc80-coated pedestals. Red and blue arrows indicate slower than normal velocity peaks, which were observed in different oscillation directions in these two examples. (**B**) Changes in the coordinate of the dumbbell beads during motor pulling or trap-induced Ndc80c sliding, tested using the same microtubule dumbbells.

Experiments with two additional Ndc80c constructs, which differed in length and subunit composition but contained wild-type microtubule-binding domains, also exhibited directional asymmetry (Fig. S8). This finding underscores that polarity-dependent sliding is an intrinsic property of the Ndc80c microtubule-binding interface, independent of its overall architecture. Asymmetric sliding persisted in experiments using densely coated Ndc80c pedestals (Fig. S9), demonstrating that this multimolecular behavior is not an emergent property but instead reflects a combined effect from asymmetric sliding of individual Ndc80c molecules. Further support for the conserved nature of frictional asymmetry comes from reports of similar behavior in ensembles of yeast Ndc80 complexes and kinetochore complexes in vitro *(37*). Moreover, Ndc80c binding to microtubules driven by kinetochore kinesin CENP-E significantly impeded their velocity *(39)*, highlighting Ndc80s’ ability to generate resistive friction under physiological, motor-driven conditions and extending its functional relevance beyond optical trapping experiments. In contrast, previous studies of the SKA complex, another microtubule-binding kinetochore protein, revealed symmetrical sliding velocities at the single-molecule level and weak retardation of CENP-E kinesin in multimolecular assays *(35, 39, 40)*. Comparing Ndc80c to other diffusing proteins with asymmetric force-velocity relationships, we found that its asymmetry parameter (1.3 nm) is substantially higher than the 0.05-0.5 nm reported for Kip3, EB1, NuMA and PRC1 proteins *(41, 42)* (Supplementary Text, Fig. S14). Together, these findings suggest that strong directional sensitivity of Ndc80c sliding is a unique and potentially critical specialization of this essential kinetochore component.

### Direction-Dependent Mobile Catch- and Slip-Bonds Drive Ndc80c’s Slip-Clutch Mechanism

The asymmetric frictional behavior of Ndc80c raises critical questions about its mechanistic origins. In the absence of external force, individual Ndc80c molecules diffuse along microtubules without directional bias. The rates of their random transitions between adjacent tubulin binding sites are characterized by the diffusion coefficient and depend on the depth of the associated potential wells, but not on the direction of transition. Consequently, during force-guided diffusion through these energy wells, sliding velocity should depend solely on well depth and not the direction of pulling. As pulling force increases, sliding velocity in both microtubule directions is expected to rise exponentially with identical initial slopes because they are dictated by the direction-independent thermal diffusion (Fig. 3A) *(41, 43, 44)*.

**Figure 3.**
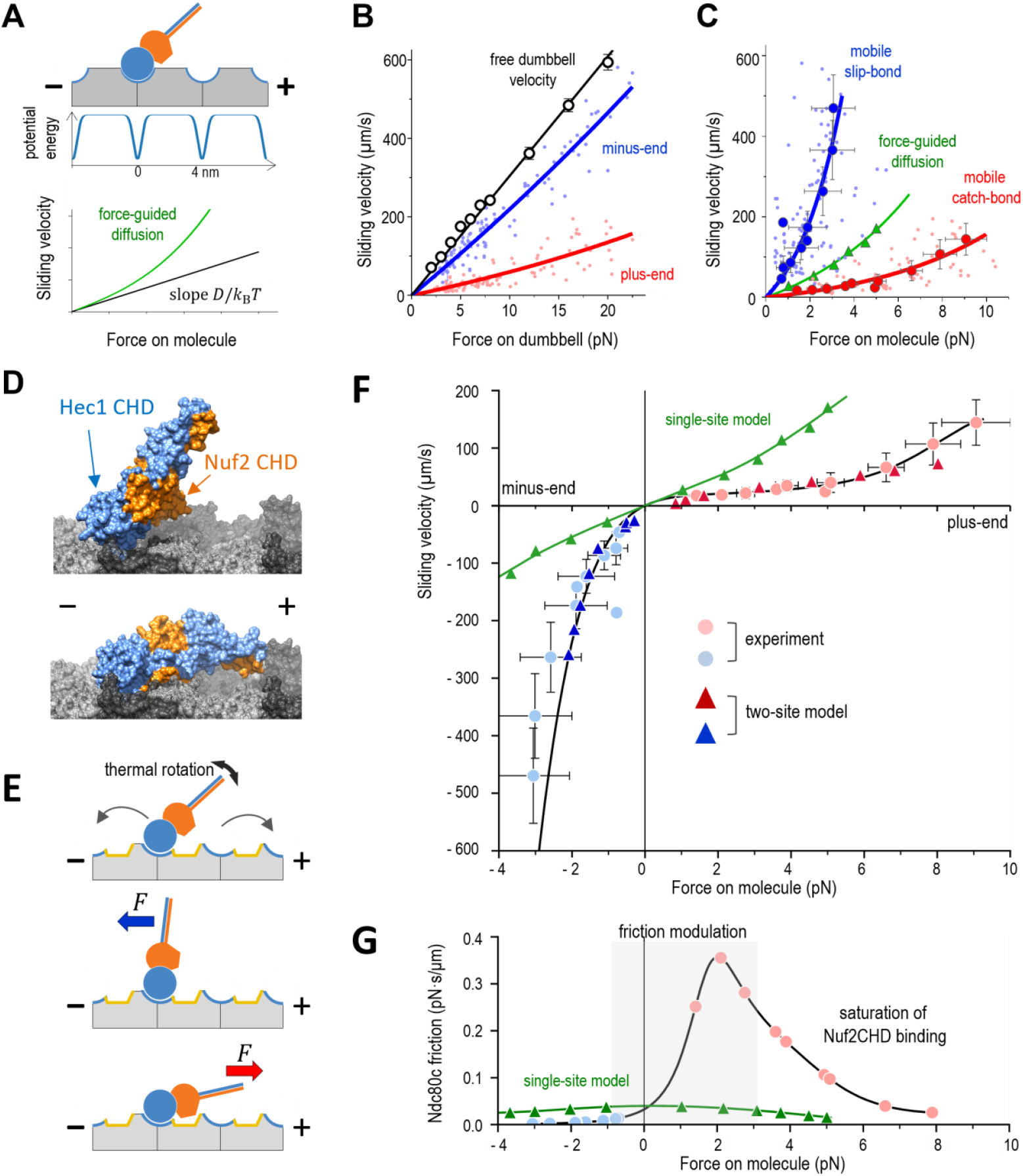
Analysis of force-velocity relationship. (**A**) Schematic of the single-site Ndc80 periodic energy landscape. The graph illustrates the expected force-velocity Bell-like relationship. At zero force, sliding velocity corresponds to the intrinsic rate of force-free diffusion: diffusion coefficient *D*, the Boltzmann’s constant *k*_*B*_, and absolute temperature *T (41)*. For the numerical details of Ndc80c relationships, refer to panel C. (**B**) Sliding velocity as a function of clamp force applied to the dumbbell, dragging Ndc80c toward the microtubule plus (red) or minus (blue) ends. Each point represents the mean Ndc80C sliding velocity during uni-directional sweeps (average 2,042 sweeps per point) from individual 30 s recordings (n = 106, N = 43 chambers). Curves are exponential fits. Open symbols show free dumbbell oscillation velocities derived from instantaneous velocity histograms. During Ndc80c sliding, lower velocity indicates stronger friction generation by Ndc80c. (**C**) Sliding velocities from panel B plotted for forces acting directly on the Ndc80c. Dark circles are binned data to represent mean ± SD and exponential approximations. Both curves deviate significantly from the theoretical prediction (green) based on the estimated force-free diffusion of 0.11 µm^2^/s, corresponding to 9 *k*_*B*_*T* well, which match closely the reported Ndc80c diffusion characteristics *(33)*. (**D**) Results of the docking computations for the crystal structures of the globular Hec1 and Nuf2 CHDs and a microtubule wall segment. The kinetochore binding end of Ndc80c and the full stalk were not modeled. (**E**) Illustration of the two-site interaction model showing microtubule-binding end of Ndc80c. Curved arrows represent 4 nm force-free diffusional steps of Ndc80 via the Hec1 CHD toe-binding. Under plus-end-directed forces, Nuf2 CHD binding frequency increases, raising the overall interaction energy. Minus-end-directed forces reduce overall binding by pulling Nuf2 CHD away from the microtubule. (**F**) Solution of the two-site Brownian model with each triangle representing the simulation with potential well depths for Hec1 and Nuf2 CHD binding sites (6 *k*_*B*_*T* each), and Ndc80c rotational stiffness (10^−2^ pN·µm). Line is polynomial fit with origin intercept. Light-colored circles show experimental data and green triangles show model results as in panel C. Negative forces correspond to minus-end-directed pulling. (**G**) Friction coefficient of Ndc80c (black line), with pale symbols indicating the force values shown in panel F. The gray area highlights the force range where the Ndc80c friction coefficient increases dramatically. The single-site model fails to capture both the magnitude and shape of this complex dependency, despite showing similar behavior at zero force.

Given the asymmetric sliding velocities of Ndc80c, we investigated which microtubule direction aligned with the expected force-velocity relationship, and which deviated from it. Using the UFFC system, we recorded bi-directional Ndc80c sliding events as brief as 2 ms under forces ranging from 2 to 20 pN, encompassing and exceeding the physiological force range experienced by microtubule-bound proteins at kinetochores *(21, 45)*. As anticipated, sliding velocities increased with higher laser trapping forces in both microtubule directions (Fig. 3B, Fig. S10). However, the resulting force-velocity relationships could not be directly interpreted because they did not represent the actual force experienced by Ndc80c. This discrepancy arises from the compliance of the microtubule dumbbell system and viscous drag, which reduce the effective force acting on the molecule. To address this, we developed a detailed mechanical model of the UFFC system to calculate the direct force experienced by sliding Ndc80c (Supplementary Materials and Methods, Figs. S11 -S12). Using this model, we replotted the sliding velocities as a function of direct molecular force. (Fig. 3C). We then compared experimental dependencies with the theoretical force-velocity relationship derived from the initial slopes of these curves. Strikingly, this analysis revealed that both sliding directions deviated from expected behavior, rather than one conforming to predictions while the other diverged (Fig. 3C).

Notably, the minus-end-directed velocity curve exhibited a significantly steeper slope than predicted, consistent with shallower potential wells for Ndc80c translocation in this direction. The reduced binding strength under force is characteristic of a “slip-bond”, leading us to refer to fast minus-end-directed translocation as “mobile slip-bond” behavior. Conversely, the plus-end-directed curve revealed slow translocation corresponding to significantly deeper energy wells, indicative of a “catch-bond” mechanism. We term this behavior “mobile catch-bond” to emphasize the dynamic nature of resistive Ndc80c sliding. Using Brownian dynamics approach, we determined that potential energy wells in the plus-end direction are ∼3 *k*_*B*_*T* deeper compared to minus-end (10-11 *k*_*B*_*T* and 7-8 *k*_*B*_*T*, respectively) (Supplementary Text, Fig. S13). This finding is remarkable, suggesting a deviation from the direction-independent principle of molecular transitions, underscoring the need for further investigation into the molecular mechanisms of this asymmetry.

### A Two-Site Model Explains Asymmetric Friction in Sliding Ndc80c

To explain the paradoxical ability of Ndc80c to generate differential friction at the single-molecule level, we evaluated several models. In ensemble measurements and sliding kinetochores, differential friction could arise from variations in the number of engaged Ndc80c molecules during plus-versus minus-end-directed motion, but this cannot account for the asymmetry observed in single-molecule experiments. Changes in Ndc80c’s overall architecture due to force were also ruled out, as the asymmetry persisted in the truncated Bonsai construct, which lacks the full structural complexity of native Ndc80c.

We hypothesized that during sliding Ndc80c engages the microtubule via two binding sites: the CHDs in the Hec1 and Nuf2 subunits. Although the Nuf2 CHD does not contact tubulin under tight molecular packing *(27)*, charge-altering mutations in this domain reduce Ndc80c microtubule-binding affinity in vitro *(25)*, suggesting that under low molecular density conditions, the Nuf2 CHD may directly engage polymerized tubulin. To explore this further, we performed in silico docking using the crystal structure of the Ndc80c’s microtubule-binding end and a microtubule wall segment. This analysis identified two low-energy binding configurations (Fig. 3D, Materials and Methods). The highest-scoring arrangement featured the Hec1 CHD as the sole binding site, with the Nuf2 CHD tilted toward the plus end, as observed in the “toe-binding” cryo-EM structure *(27)*. The second-best configuration, resembling “belly’ binding”, revealed an extended interface involving both CHDs, with the Nuf2 CHD aligned toward the plus end. While docking does not definitively establish this binding mode, these findings, combined with molecular-genetic evidence, suggest that force modulates Nuf2 CHD engagement.

Specifically, we hypothesize that plus-end-directed force induces a conformational shift in Ndc80c, such as bending or tilting, bringing the Nuf2 CHD into direct contact with tubulin and expanding the interaction footprint (Fig. 3E). To test this, we developed a Brownian dynamics model in which Ndc80c, represented as a rod with two microtubule-binding points, tilts in a force-dependent manner (Fig. S15, Supplementary Text). Simulations applying force to the distal end of the rod revealed a naturally emergent asymmetric force-velocity relationship that closely matches experimental data under realistic parameters (Fig. 3F, Video 1). Unlike the single-site model with Hec1 CHD only, which fits experimental force-velocity with distinct dependencies for each direction, the two-site model produces a continuous force-velocity function that satisfies the thermodynamic constraint of a direction-independent initial slope—providing the first physically realistic description of asymmetric sliding in any diffusing protein (Figs. S16-S17, Supplementary Text).

We derived the force–friction relationship for sliding Ndc80c from the slope of the experimental force– velocity curve. Ndc80c generates little resistance to minus-end-directed translocation, with friction almost insensitive to molecular force (Fig. 3G). However, the friction coefficient increases sharply from –1 pN to + 2-3 pN force, overlapping with the thermally-dominated regime. As plus-end-directed force increases, friction declines once Nuf2 CHD engagement nears saturation, following the expected monotonically decreasing pattern for force-guided diffusion. In contrast, the model where Ndc80c binds solely via the Hec1 CHD predicts a peak in friction at zero force and a slight, symmetrical decline, consistent with a conventional force–velocity relationship. The >10-fold friction range observed for Ndc80c is incompatible with the single-site model and represents a unique adaptation for force-modulated single-molecule operation near thermal boundaries.

### Nuf2 CHD modulates molecular friction in sliding Ndc80c

According to the two-site model, the CHDs of both the Hec1 and Nuf2 subunits govern the sliding behavior of Ndc80c, with the Hec1 CHD anchoring Ndc80c to the microtubule wall and the Nuf2 CHD regulating molecular friction. Given that the unstructured Hec1 tail lacks a defined tubulin interaction footprint, it is expected to contribute less to molecular friction (Fig. 4A). To test these predictions, we first examined the force-dependent sliding of Ndc80c harboring a charge-reversing mutation, K166D, in the toe region of the Hec1 CHD, which severely disrupts microtubule binding in the absence of force *(25)*. Under dragging force, this mutant exhibited significantly faster plus-end-directed sliding, consistent with the Hec1 CHD’s acting as the primary microtubule-binding site (Fig. 4 B,C). In contrast, deletion of the Hec1 tail, despite dramatically reducing microtubule-binding affinity *(26, 33)*, preserved strong friction, as reflected in its slow plus-end sliding velocity (Fig. 4 B-E). These findings indicate that reduced binding affinity under no-force conditions does not necessarily impair frictional properties in the sliding Ndc80c clamp. In cells, phosphorylation of the Hec1 tail modulates resistive friction on polymerizing microtubules, but its effects are far weaker than perturbations in the Hec1 CHD itself, aligning with our observations *(23, 25, 28, 38, 46, 47)*.

**Figure 4.**
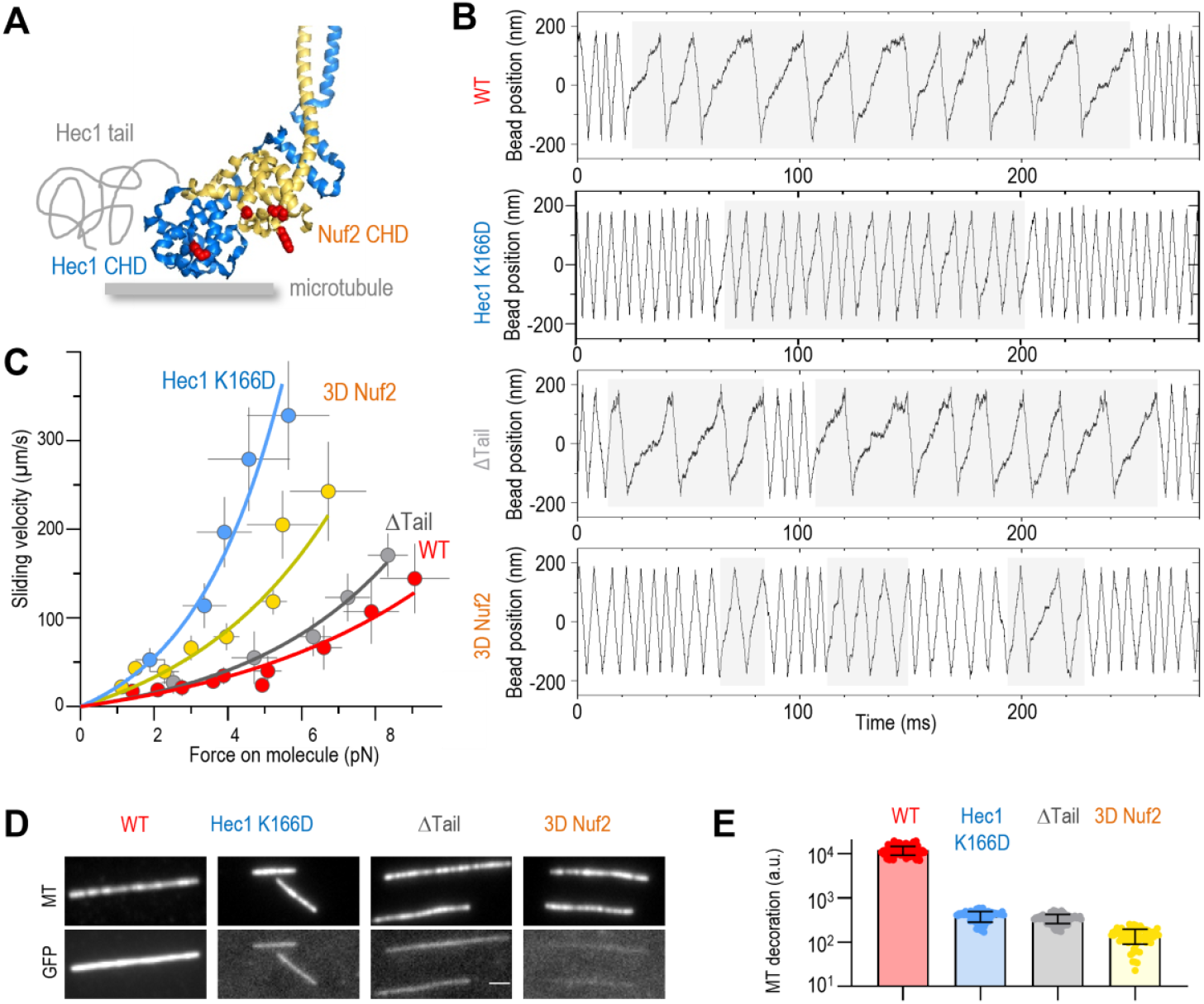
Nuf2 CHD, but not Hec1 tail, is required for friction generation. (**A**) Crystal structure of the Hec1 and Nuf2 CHDs (PDB# 2VE7) *(25)* showing mutated residues (red) and the not-to-scale drawing of the unstructured Hec1 N-terminal extension. (**B**) Example bead coordinate signals in experiments using the indicated Ndc80c Bonsai proteins at 4 pN clamp force. Tandem slow velocity segments (highlighted with light gray) are evident for all proteins, but gliding is faster in CHDs mutants. (**C**) Force-velocity dependencies for the plus-end-directed sliding determined with UFFC. Each symbol shows mean and SD for different force bins based on N = 43 chambers for WT, N = 6 for ΔTail; N = 12 for Nuf2 3D; N = 8 for Hec1 K166D. (**D**) Images of rhodamine-labeled taxol-stabilized microtubules and their decoration by the indicated GFP-tagged Ndc80c Bonsai proteins (100 nM). Image with unmodified Bonsai Ndc80c-GFP has reduced contrast relative to other GFP images because of its excessive brightness. (**E**) Brightness of microtubule decoration with indicated GFP-tagged Ndc80c proteins plotted on the semi-log scale. Each dot shows average GFP fluorescence of one microtubule, bars and whiskers show mean ± SD.

We next tested the predicted contribution of the Nuf2 CHD to friction generation. To this end, we used an Ndc80c construct carrying triple charge-reversing mutations in the positively charged face of the Nuf2 CHD (3D Nuf2: K33D, K41D, K115D) (Fig. 4A). The 3D Nuf2 mutant exhibited significantly reduced microtubule-wall affinity but retained its ability to slide under force in the UFFC assay. If the Nuf2 CHD functioned solely as a binding hub for other kinetochore proteins, these mutations would not be expected to alter Ndc80c sliding friction. However, consistent with our model, the 3D Nuf2 mutant exhibited significantly faster plus-end-directed sliding compared to wild-type Ndc80c (Fig. 4 B,C), with an estimated maximum friction coefficient fivefold lower than that of wild-type Ndc80c. These results demonstrate that the Nuf2 CHD plays a direct role in the sliding clutch mechanism by modulating molecular friction with microtubules rather than acting solely as a passive scaffold. Furthermore, they suggest a molecular basis for Nuf2’s role in generating tension across sister kinetochores in mitotic cells *(28)*.

## Discussion

### Implications for kinetochore coupling mechanisms

The exceptional temporal resolution of the UFFC assay makes it ideally suited for studying the force-dependent motility of low-affinity proteins like Ndc80c at the single-molecule level. Unlike techniques that drag protein-coated beads at constant velocity along stationary microtubules, the UFFC minimizes potential artifacts, such as slippage caused by bead rolling *(48)*, by applying lateral forces to molecules sliding at their natural, force-dependent velocities. Using this powerful approach, combined with mechanistic modeling, we uncovered a novel mechanism by which a non-motor protein translocates at rates differing by more than an order of magnitude depending on microtubule polarity, representing a sliding molecular clutch.

Molecular clutches, described in various cytoskeletal processes, dynamically engage and disengage molecular interfaces in response to mechanical forces, modulating bond lifetime *(49-51)*. The Ndc80 sliding clutch, however, represents a fundamental departure from prior models—rather than modulating duration of static bonds, it controls the velocity of directional motion under force at the single-molecule level. This dual mobile catch- and slip-bond behavior, while non-intuitive, aligns with thermodynamic principles and arises naturally from force-induced conformational shift of Ndc80c. The Hec1 CHD provides the primary microtubule contact, anchoring Ndc80c and enabling force transmission, while the Nuf2 CHD modulates molecular friction in a direction-dependent manner, revealing an unrecognized role for this domain as a friction-regulator. Notably, friction modulation operates near thermal limits, remaining effective despite thermal fluctuations and within physiological force ranges, as single microtubule-bound proteins experience forces below 5-6 pN, the estimated maximal load exerted by a depolymerizing tubulin protofilament *(52)*.

Prior in vitro studies reveal that thermal diffusion and microtubule-wall affinity of Ndc80c do not stand out among other kinetochore proteins *(39)*, leading to a long-standing puzzle: what makes it uniquely suited for microtubule-end coupling in cells? Interestingly, multimolecular assemblies of other microtubule-associated proteins, such as the SKA complex, CLASP2, EB1 proteins, and the microtubule-binding tail of CENP-E kinesin, generate far less molecular friction than Ndc80c *(39)*. While small frictional asymmetries have been reported for some spindle-associated proteins *(42)*, single-molecule studies reveal symmetric friction in kinetochore-associated SKA complexes *(35)*. Thus, the sliding clutch behavior provides a compelling explanation for Ndc80’s unique importance in kinetochore–microtubule coupling.

We propose that at kinetochores, sliding Ndc80c resists forces pulling the kinetochore away from the microtubule plus-end, thereby guarding against end-detachment, while under pole-directed forces, it forms highly mobile bonds that minimize counterproductive drag and assist motion away from the end. This sliding-clutch functions not as a binary, direction-dependent frictional switch but as a continuously adjustable system operating at the nanosecond scale. A dynamic modulation is essential because, even under constant external loads, thermal noise introduces force fluctuations, necessitating rapid Ndc80c adaptation to maintain efficient sliding. Such a mechanism is ideally suited for kinetochore-microtubule coupling across diverse motility modes and mitotic stages. In cells, additional factors likely adjust kinetochore slip-clutch behavior, including distinct protein activities at opposite kinetochores, variations in the number of friction-generating proteins and their post-translational modifications, and force-induced changes in kinetochore architecture or microtubule bundle composition *(18, 37, 46, 53-60)*. Investigating the roles of these factors and their unique contributions presents a significant and ongoing challenge for future research.

These molecular findings about Ndc80c fundamentally advance our understanding of kinetochore motility, providing a new biophysical framework for end-coupling *(21)*. In the prevailing “biased diffusion” model, an ensemble of mechanically linked Ndc80c molecules exhibits direction-independent friction *(19, 20, 61)*. End-coupling results from a slow diffusional search for the minimal energy configuration under force, which gradually pulls the linked molecular ensemble beyond the microtubule end, forming and overhang. Our findings challenge relevance of this concept, demonstrating that expedient frictional modulation occurs at the level of individual wall-bound Ndc80 molecules, calling for new physical models of kinetochore coupling mechanisms. Intriguingly, recent studies highlight the Nuf2 subunit as a pivotal molecular interface for regulatory kinases and other microtubule-binding proteins *(29-31)*. The direct interaction between tension-sensitive Ndc80c conformations and mitotic regulators may provide a mechanism for integrating mechanical cues with mitotic signaling, offering an exciting frontier for future research.

## Acknowledgements

We thank Dr. F Balabin for preparing computational video and M. Godzi for molecular docking. We are grateful to Dr. J. DeLuca (Colorado State Univ. at Ft. Collins) for providing purified Ndc80 mutant proteins and Ndc80 Bronsai, and to Dr. E. Holzbaur (Univ. of Pennsylvania) for purified dynein. We thank Drs. I. Cheeseman, J.R. McIntosh and J. DeLuca for critical reading and feedback on the manuscript; Dr. S. Pyrpassopoulos for helpful tips on three bead assay; Drs. M. Ostap and Y. Goldman for insightful discussions; to Dr. E. Tarasovetc, P.-T. Chen, V. Mustyatsa and Dr. L. Wangxi for help with protein purification, A. Maiorov and other members of Grishchuk lab for general assistance and discussions.

## Author Contributions

V.M.D. and S.K.T. performed experiments, F.I.A. and V.M.D. constructed UFFC instrument, I.V.G. and F.I.A. carried out theoretical modeling, E.L.G. designed the research project and wrote the paper with input from all co-authors.

## Data and materials availability

All data needed to evaluate the conclusions in the paper are available in the main text, Supplementary Materials, and Data Source file.

## Supplementary materials

Materials and Methods

Supplementary Text

Figs. S1 to S18

Tables S1 to S5

Movie S1

Data Source file S1

## References

1. C. E. Walczak, S. Cai, A. Khodjakov, Mechanisms of chromosome behaviour during mitosis. Nature Reviews Molecular Cell Biology 11, 91–102 (2010).

2. E. Vladimirou, E. Harry, N. Burroughs, A. D. McAinsh, Springs, clutches and motors: driving forward kinetochore mechanism by modelling. Chromosome Research 19, 409–421 (2011).

3. J. K. Monda, I. M. Cheeseman, The kinetochore–microtubule interface at a glance. Journal of Cell Science 131, (2018).

4. R. Skibbens, V. Skeen, E. Salmon, Directional instability of kinetochore motility during chromosome congression and segregation in mitotic newt lung cells: a push-pull mechanism. Journal of Cell Biology 122, 859–875 (1993).

5. R. V. Skibbens, C. L. Rieder, E. D. Salmon, Kinetochore motility after severing between sister centromeres using laser microsurgery: Evidence that kinetochore directional instability and position is regulated by tension. Journal of Cell Science 108, 2537–2548 (1995).

6. A. Khodjakov, C. L. Rieder, Kinetochores moving away from their associated pole do not exert a significant pushing force on the chromosome. Journal of Cell Biology 135, 315–327 (1996).

7. R.-H. Chen, J. C. Waters, E. D. Salmon, A. W. Murray, Association of Spindle Assembly Checkpoint Component XMAD2 with Unattached Kinetochores. Science 274, 242–246 (1996).

8. S. Dumont, T. J. Mitchison, Force and Length in the Mitotic Spindle. Current Biology 19, R749–R761 (2009).

9. S. Dumont, E. D. Salmon, T. J. Mitchison, Deformations Within Moving Kinetochores Reveal Different Sites of Active and Passive Force Generation. Science 337, 355–358 (2012).

10. P. Maddox, A. Straight, P. Coughlin, T. J. Mitchison, E. D. Salmon Direct observation of microtubule dynamics at kinetochores in Xenopus extract spindles : implications for spindle mechanics. Journal of Cell Biology 162, 377–382 (2003).

11. R. B. Nicklas, Measurements of the force produced by the mitotic spindle in anaphase. J. Cell Biol. 97, 542–548 (1983).

12. E. L. Grishchuk, M. I. Molodtsov, F. I. Ataullakhanov, J. R. McIntosh, Force production by disassembling microtubules. Nature 438, 384–388 (2005).

13. A. Efremov, E. L. Grishchuk, J. R. McIntosh, F. I. Ataullakhanov, In search of an optimal ring to couple microtubule depolymerization to processive chromosome motions. Proceedings of the National Academy of Sciences 104, 19017–19022 (2007).

14. A. Musacchio, A. Desai, A Molecular View of Kinetochore Assembly and Function. Biology 6, 5 (2017).

15. I. M. Cheeseman, S. Anderson, M. Jwa, E. M. Green, J.-s. Kang, J. R. Yates, III, C. S. M. Chan, D. G. Drubin, G. Barnes, Phospho-Regulation of Kinetochore-Microtubule Attachments by the Aurora Kinase Ipl1p. Cell 111, 163–172 (2002).

16. J. G. DeLuca, W. E. Gall, C. Ciferri, D. Cimini, A. Musacchio, E. D. Salmon, Kinetochore Microtubule Dynamics and Attachment Stability Are Regulated by Hec1. Cell 127, 969–982 (2006).

17. A. V. Zaytsev, L. J. R. Sundin, K. F. DeLuca, E. L. Grishchuk, J. G. DeLuca, Accurate phosphoregulation of kinetochore–microtubule affinity requires unconstrained molecular interactions. Journal of Cell Biology 206, 45–59 (2014).

18. A. A. Kukreja, S. Kavuri, A. P. Joglekar, Microtubule Attachment and Centromeric Tension Shape the Protein Architecture of the Human Kinetochore. Current Biology 30, 4869–4881.e4865 (2020).

19. T. L. Hill, Theoretical problems related to the attachment of microtubules to kinetochores. Proceedings of the National Academy of Sciences 82, 4404–4408 (1985).

20. A. F. Powers, A. D. Franck, D. R. Gestaut, J. Cooper, B. Gracyzk, R. R. Wei, L. Wordeman, T. N. Davis, C. L. Asbury, The Ndc80 Kinetochore Complex Forms Load-Bearing Attachments to Dynamic Microtubule Tips via Biased Diffusion. Cell 136, 865–875 (2009).

21. E. L. Grishchuk, Biophysics of Microtubule End Coupling at the Kinetochore in Centromeres and Kinetochores: Discovering the Molecular Mechanisms Underlying Chromosome Inheritance, B. E. Black, Ed. (Springer International Publishing, Cham, 2017), vol. 56, pp. 397–428.

22. J. G. Tooley, S. A. Miller, P. T. Stukenberg, The Ndc80 complex uses a tripartite attachment point to couple microtubule depolymerization to chromosome movement. Molecular Biology of the Cell 22, 1217–1226 (2011).

23. R. T. Wimbish, J. G. DeLuca, Hec1/Ndc80 Tail Domain Function at the Kinetochore-Microtubule Interface. Frontiers in Cell and Developmental Biology 8, (2020).

24. G. M. Alushin, V. Musinipally, D. Matson, J. Tooley, P. T. Stukenberg, E. Nogales, Multimodal microtubule binding by the Ndc80 kinetochore complex. Nature Structural & Molecular Biology 19, 1161–1167 (2012).

25. C. Ciferri, S. Pasqualato, E. Screpanti, G. Varetti, S. Santaguida, G. Dos Reis, A. Maiolica, J. Polka, J. G. De Luca, P. De Wulf, M. Salek, J. Rappsilber, C. A. Moores, E. D. Salmon, A. Musacchio, Implications for Kinetochore-Microtubule Attachment from the Structure of an Engineered Ndc80 Complex. Cell 133, 427–439 (2008).

26. R. R. Wei, J. Al-Bassam, S. C. Harrison, The Ndc80/HEC1 complex is a contact point for kinetochore-microtubule attachment. Nature Structural & Molecular Biology 14, 54–59 (2007).

27. G. M. Alushin, V. H. Ramey, S. Pasqualato, D. A. Ball, N. Grigorieff, A. Musacchio, E. Nogales, The Ndc80 kinetochore complex forms oligomeric arrays along microtubules. Nature 467, 805–810 (2010).

28. L. J. R. Sundin, G. J. Guimaraes, J. G. DeLuca, The NDC80 complex proteins Nuf2 and Hec1 make distinct contributions to kinetochore–microtubule attachment in mitosis. Molecular Biology of the Cell 22, 759–768 (2011).

29. J. A. Zahm, S. C. Harrison, A communication hub for phosphoregulation of kinetochore-microtubule attachment. Current Biology 34, 2308–2318.e2306 (2024).

30. R. Pleuger, C. Cozma, S. Hohoff, C. Denkhaus, A. Dudziak, F. Kaschani, M. Kaiser, A. Musacchio, I. R. Vetter, S. Westermann, Microtubule end-on attachment maturation regulates Mps1 association with its kinetochore receptor. Current Biology 34, 2279–2293.e2276 (2024).

31. E. J. Parnell, E. E. Jenson, M. P. Miller, A conserved site on Ndc80 complex facilitates dynamic recruitment of Mps1 to yeast kinetochores to promote accurate chromosome segregation. Current Biology 34, 2294–2307.e2294 (2024).

32. A. F. Powers, A. D. Franck, D. R. Gestaut, J. Cooper, B. Gracyzk, R. R. Wei, L. Wordeman, T. N. Davis, C. L. Asbury, The Ndc80 kinetochore complex uses biased diffusion to couple chromosomes to dynamic microtubule tips. Cell 136, 865–875 (2009).

33. A. V. Zaytsev, J. E. Mick, E. Maslennikov, B. Nikashin, J. G. DeLuca, E. L. Grishchuk, Multisite phosphorylation of the NDC80 complex gradually tunes its microtubule-binding affinity. Mol. Biol. Cell 26, 1829–1844 (2015).

34. M. Capitanio, M. Canepari, M. Maffei, D. Beneventi, C. Monico, F. Vanzi, R. Bottinelli, F. S. Pavone, Ultrafast force-clamp spectroscopy of single molecules reveals load dependence of myosin working stroke. Nat Methods 9, 1013–1019 (2012).

35. S. K. Tripathy, V. M. Demidov, I. V. Gonchar, S. Wu, F. I. Ataullakhanov, E. L. Grishchuk, Ultrafast Force-Clamp Spectroscopy of Microtubule-Binding Proteins in Optical Tweezers: Methods and Protocols, Methods in Molecular Biology, vol. 2478, A. Gennerich, Ed. (Springer, 2022), pp. 609–650.

36. M. S. Woody, D. A. Winkelmann, M. Capitanio, E. M. Ostap, Y. E. Goldman, Single molecule mechanics resolves the earliest events in force generation by cardiac myosin. eLife 8, e49266 (2019).

37. J. D. Larson, N. A. Heitkamp, L. E. Murray, A. R. Popchock, S. Biggins, C. L. Asbury, Kinetochores grip microtubules with directionally asymmetric strength. Journal of Cell Biology 224, (2024).

38. A. Suzuki, B. L. Badger, J. Haase, T. Ohashi, H. P. Erickson, E. D. Salmon, K. Bloom, How the kinetochore couples microtubule force and centromere stretch to move chromosomes. Nature Cell Biology 18, 382–392 (2016).

39. M. Chakraborty, E. V. Tarasovetc, A. V. Zaytsev, M. Godzi, A. C. Figueiredo, F. I. Ataullakhanov, E. L. Grishchuk, Microtubule end conversion mediated by motors and diffusing proteins with no intrinsic microtubule end-binding activity. Nat. Commun. 10, 1673–1687 (2019).

40. J. K. Monda, I. P. Whitney, E. V. Tarasovetc, E. Wilson-Kubalek, R. A. Milligan, E. L. Grishchuk, I. M. Cheeseman, Microtubule Tip Tracking by the Spindle and Kinetochore Protein Ska1 Requires Diverse Tubulin-Interacting Surfaces. Current Biology 27, 3666–3675.e3666 (2017).

41. V. Bormuth, V. Varga, J. Howard, E. Schäffer, Protein Friction Limits Diffusive and Directed Movements of Kinesin Motors on Microtubules. Science 325, 870–873 (2009).

42. S. Forth, K.-C. Hsia, Y. Shimamoto, Tarun M. Kapoor, Asymmetric Friction of Nonmotor MAPs Can Lead to Their Directional Motion in Active Microtubule Networks. Cell 157, 420–432 (2014).

43. G. I. Bell, Models for the Specific Adhesion of Cells to Cells. Science 200, 618–627 (1978).

44. H. Suda, Origin of Friction Derived from Rupture Dynamics. Langmuir 17, 6045–6047 (2001).

45. A. A. Ye, S. Cane, T. J. Maresca, Chromosome biorientation produces hundreds of piconewtons at a metazoan kinetochore. Nat. Commun. 7, 1–9 (2016).

46. A. F. Long, D. B. Udy, S. Dumont, Hec1 Tail Phosphorylation Differentially Regulates Mammalian Kinetochore Coupling to Polymerizing and Depolymerizing Microtubules. Current Biology 27, 1692–1699.e1693 (2017).

47. R. T. Wimbish, K. F. DeLuca, J. E. Mick, J. Himes, I. Jiménez-Sánchez, A. A. Jeyaprakash, J. G. DeLuca, The Hec1/Ndc80 tail domain is required for force generation at kinetochores, but is dispensable for kinetochore-microtubule attachment formation and Ska complex recruitment. Mol Biol Cell 31, 1453–1473 (2020).

48. E. L. Grishchuk, I. S. Spiridonov, V. A. Volkov, A. Efremov, S. Westermann, D. Drubin, G. Barnes, F. I. Ataullakhanov, J. R. McIntosh, Different assemblies of the DAM1 complex follow shortening microtubules by distinct mechanisms. Proceedings of the National Academy of Sciences 105, 6918–6923 (2008).

49. G. Giannone, R.-M. Mège, O. Thoumine, Multi-level molecular clutches in motile cell processes. Trends in Cell Biology 19, 475–486 (2009).

50. F. B. Cleary, M. A. Dewitt, T. Bilyard, Z. M. Htet, V. Belyy, D. D. Chan, A. Y. Chang, A. Yildiz, Tension on the linker gates the ATP-dependent release of dynein from microtubules. Nature Communications 5, 4587 (2014).

51. D. L. Huang, N. A. Bax, C. D. Buckley, W. I. Weis, A. R. Dunn, Vinculin forms a directionally asymmetric catch bond with F-actin. Science 357, 703–706 (2017).

52. M. I. Molodtsov, E. L. Grishchuk, A. K. Efremov, J. R. McIntosh, F. I. Ataullakhanov, Force production by depolymerizing microtubules: A theoretical study. Proceedings of the National Academy of Sciences 102, 4353–4358 (2005).

53. J. S. Tirnauer, J. C. Canman, E. D. Salmon, T. J. Mitchison, EB1 Targets to Kinetochores with Attached, Polymerizing Microtubules. Molecular Biology of the Cell 13, 4308–4316 (2002).

54. A. C. Amaro, C. P. Samora, R. Holtackers, E. Wang, I. J. Kingston, M. Alonso, M. Lampson, A. D. McAinsh, P. Meraldi, Molecular control of kinetochore-microtubule dynamics and chromosome oscillations. Nature Cell Biology 12, 319–329 (2010).

55. M. Rosas-Salvans, C. Rux, M. Das, S. Dumont, SKAP binding to microtubules reduces friction at the kinetochore-microtubule interface and increases attachment stability under force. bioRxiv, (2024).

56. M. Rosas-Salvans, R. Sutanto, P. Suresh, S. Dumont, The Astrin-SKAP complex reduces friction at the kinetochore-microtubule interface. Current Biology 32, 2621–2631.e2623 (2022).

57. C. Castrogiovanni, A. V. Inchingolo, J. U. Harrison, D. Dudka, O. Sen, N. J. Burroughs, A. D. McAinsh, P. Meraldi, Evidence for a HURP/EB free mixed-nucleotide zone in kinetochore-microtubules. Nature Communications 13, 4704 (2022).

58. J. P. I. Welburn, M. Vleugel, D. Liu, J. R. Yates, III, M. A. Lampson, T. Fukagawa, I. M. Cheeseman, Aurora B Phosphorylates Spatially Distinct Targets to Differentially Regulate the Kinetochore-Microtubule Interface. Molecular Cell 38, 383–392 (2010).

59. E. Roscioli, T. E. Germanova, C. A. Smith, P. A. Embacher, M. Erent, A. I. Thompson, N. J. Burroughs, A. D. McAinsh, Ensemble-Level Organization of Human Kinetochores and Evidence for Distinct Tension and Attachment Sensors. Cell Reports 31, (2020).

60. T. Y. Yoo, J.-M. Choi, W. Conway, C.-H. Yu, R. V. Pappu, D. J. Needleman, Measuring NDC80 binding reveals the molecular basis of tension-dependent kinetochore-microtubule attachments. eLife 7, e36392 (2018).

61. A. P. Joglekar, A. J. Hunt, A Simple, Mechanistic Model for Directional Instability during Mitotic Chromosome Movements. Biophysical Journal 83, 42–58 (2002).

62. H. P. Miller, L. Wilson, Preparation of microtubule protein and purified tubulin from bovine brain by cycles of assembly and disassembly and phosphocellulose chromatography in Methods Cell Biol. (Elsevier, 2010), vol. 95, pp. 2–15.

63. A. Hyman, D. Drechsel, D. Kellogg, S. Salser, K. Sawin, P. Steffen, L. Wordeman, T. Mitchison, Preparation of modified tubulins in Methods Enzymol. (Elsevier, 1991), vol. 196, pp. 478–485.

64. J. C. Schmidt, H. Arthanari, A. Boeszoermenyi, N. M. Dashkevich, E. M. Wilson-Kubalek, N. Monnier, M. Markus, M. Oberer, R. A. Milligan, M. Bathe, G. Wagner, E. L. Grishchuk, I. M. Cheeseman, The kinetochore-bound Ska1 complex tracks depolymerizing microtubules and binds to curved protofilaments. Dev. Cell 23, 968–980 (2012).

65. R. B. Case, D. W. Pierce, N. HomBooher, C. L. Hart, R. D. Vale, The directional preference of kinesin motors is specified by an element outside of the motor catalytic domain. Cell 90, 959–966 (1997).

66. S. Toba, Y. Y. Toyoshima, Dissociation of double-headed cytoplasmic dynein into single-headed species and its motile properties. Cell Motil Cytoskel 58, 281–289 (2004).

67. N. Gudimchuk, E. V. Tarasovetc, V. Mustyatsa, A. L. Drobyshev, B. Vitre, D. W. Cleveland, F. I. Ataullakhanov, E. L. Grishchuk, Probing mitotic CENP-E kinesin with the tethered cargo motion assay and laser tweezers. Biophys. J. 114, 2640–2652 (2018).

68. M. Chakraborty, E. V. Tarasovetc, E. L. Grishchuk, In vitro reconstitution of lateral to end-on conversion of kinetochore–microtubule attachments in Methods Cell Biol. (Elsevier, 2018), vol. 144, pp. 307–327.

69. V. A. Volkov, A. V. Zaytsev, E. L. Grishchuk, Preparation of segmented microtubules to study motions driven by the disassembling microtubule ends. J Vis. Exp., e51150 (2014).

70. M. Barisic, R. Silva e Sousa, S. K. Tripathy, M. M. Magiera, A. V. Zaytsev, A. L. Pereira, C. Janke, E. L. Grishchuk, H. Maiato, Microtubule detyrosination guides chromosomes during mitosis. Science 348, 799–803 (2015).

71. D. Kozakov, D. R. Hall, B. Xia, K. A. Porter, D. Padhorny, C. Yueh, D. Beglov, S. Vajda, The ClusPro web server for protein-protein docking. Nat Protoc 12, 255–278 (2017).

72. L. Gardini, A. Tempestini, F. S. Pavone, M. Capitanio, High-Speed Optical Tweezers for the Study of Single Molecular Motors in Molecular Motors: Methods and Protocols, C. Lavelle, Ed. (Springer New York, New York, NY, 2018), pp. 151–184.

73. J. Gao, W. D. Luedtke, D. Gourdon, M. Ruths, J. N. Israelachvili, U. Landman, Frictional Forces and Amontons’ Law: From the Molecular to the Macroscopic Scale. The Journal of Physical Chemistry B 108, 3410–3425 (2004).

74. P. Hänggi, P. Talkner, M. Borkovec, Reaction-rate theory: fifty years after Kramers. Reviews of Modern Physics 62, 251–341 (1990).

75. P. T. Boggs, J. R. Donaldson. (, National Institute of Standards and Technology, Gaithersburg, MD, 1989).

76. D. E. Dupuis, W. H. Guilford, J. Wu, D. M. Warshaw, Actin filament mechanics in the laser trap. Journal of Muscle Research & Cell Motility 18, 17–30 (1997).

77. C. Veigel, C. F. Schmidt, Moving into the cell: single-molecule studies of molecular motors in complex environments. Nature Reviews Molecular Cell Biology 12, 163–176 (2011).

78. Y. Takagi, E. E. Homsher, Y. E. Goldman, H. Shuman, Force generation in single conventional actomyosin complexes under high dynamic load. Biophys J 90, 1295–1307 (2006).

79. J. Happel, H. Brenner, Wall Effects on the Motion of a Single Particle in Low Reynolds number hydrodynamics: with special applications to particulate media. (Springer Science & Business Media, 2012), vol. 1, pp. 286–357.

80. Z. Lansky, M. Braun, A. Lüdecke, M. Schlierf, Pieter R. ten Wolde, Marcel E. Janson, S. Diez, Diffusible Crosslinkers Generate Directed Forces in Microtubule Networks. Cell 160, 1159–1168 (2015).

81. G. Arpağ, S. Shastry, William O. Hancock, E. Tüzel, Transport by Populations of Fast and Slow Kinesins Uncovers Novel Family-Dependent Motor Characteristics Important for In Vivo Function. Biophysical Journal 107, 1896–1904 (2014).

82. I. M. Cheeseman, J. S. Chappie, E. M. Wilson-Kubalek, A. Desai, The Conserved KMN Network Constitutes the Core Microtubule-Binding Site of the Kinetochore. Cell 127, 983–997 (2006).

83. E. M. Wilson-Kubalek, I. M. Cheeseman, C. Yoshioka, A. Desai, R. A. Milligan Orientation and structure of the Ndc80 complex on the microtubule lattice. Journal of Cell Biology 182, 1055–1061 (2008).

84. M. M. Tirado, C. L. Martínez, J. G. de la Torre, Comparison of theories for the translational and rotational diffusion coefficients of rod-like macromolecules. Application to short DNA fragments. The Journal of Chemical Physics 81, 2047–2052 (1984).

