## Supplemental Materials for "Ndc80 complex, a conserved coupler for kinetochore-microtubule motility, is a sliding molecular clutch"

†: equal contribution

**This PDF file includes:**

Materials and Methods
- Experimental procedures
- Analysis of the UFFC recordings

Supplementary Text
- Theoretical modelling

Supplementary Tables
Supplementary Video legend
Supplementary Figures with legends
References

**Abbreviations**: BG – benzylguanine, DIG – digoxigenin, GBP – GFP-binding protein, PEG - polyethylene glycol, QPD – quadrant photodetector, SD – standard deviation, UFFC – ultrafast force-clamp

### MATERIALS AND METHODS

#### Experimental procedures

Proteins and reagents

All chemicals and reagents were purchased from Sigma (St. Louis, MO), unless specified otherwise. Tubulin was purified from cow brains by thermal cycling and chromatography, as in *(62)*. Labeling of tubulin with rhodamine (ThermoFisher, cat. # C1171) or digoxigenin (DIG) (ThermoFisher, cat. # A2952) was carried out as in *(63)*; the labeled and label-free tubulin was cycled twice to ensure that it is highly competent. Other proteins were expressed in E. coli and purified using published protocols: human Ndc80c ("Bonsai" Ndc80-GFP referred to as WT *(33)*, human "Broccoli" Ndc80-GFP *(64)*. Mutant Ndc80c Bonsai proteins (ΔTail with a deletion of 80 amino acids at the N-terminus of Hec1 chain, Hec1 K166D and 3D Nuf2 containing K33D, K41D and K115D substitutions in the Nuf2 subunit) and Human "Bronsai" Ndc80-GFP were a gift from J. DeLuca, University of Colorado. Truncated kinesin-1 (K560-GFP) was purified as in *(65)*. Cow brain dynein purified as in *(66)* was a gift from E. Holzbaur; purified GFP was kindly provided by I. Cheeseman. The adaptor protein with SNAP-tag (New England Biolabs) and GFP-binding protein (GBP) was constructed and purified as in *(67)*. Prior to each experiment, a thawed protein aliquot was centrifuged to remove aggregates and soluble concentration of GFP-labeled protein was measured via fluorescence intensity, as in *(68)*. Microtubules were polymerized using a mixture of bovine tubulins: 7.1 mg/ml unlabeled, 0.9 mg/ml DIG-labeled and 0.3 mg/ml rhodamine-labeled. The mixture was incubated at 37 °C for 25 min and stabilized with 10 µM taxol (Sigma, cat. # T7402), as in *(68)*. Microtubules were kept at room temperature in the dark for not longer than 3 days.

Preparation of the dumbbell beads

We employed two bead preparation methods to form robust dumbbells with DIG-labeled microtubules *(35)*. In the first method, streptavidin-coated polystyrene beads 0.54 µm diameter (Spherotech, cat. # SVP-05-10) were incubated with 0.11 mg/ml biotinylated anti-sheep antibodies (Jackson ImmunoResearch, cat. # 313-065-003). Beads were blocked with 1 mM biotinylated polyethylene glycol (PEG) (Quanta Biodesign, cat. #721431-18-1) and subsequently incubated with 0.01 mg/ml of sheep anti-DIG antibodies (Roche, cat. # 11333089001). For the second method, carboxylated polystyrene beads 0.51 µm diameter (Spherotech, cat. # CP-05-10) were activated in MES-Tween buffer (25 mM MES pH 5.0, 0.05% Tween20 (Sigma, cat. # P1379-25)) using water-soluble 1-ethyl-3-[3-dimethylaminopropyl] carbodiimide (Sigma, cat. # 22980) and N-hydroxysulfosuccinimide (Sigma, cat. # 56845), followed by incubation with 0.17 mg/ml anti-DIG Fab antibody fragments (Roche, cat. # 11214667001). Microtubule dumbbells formed using these bead preparation methods exhibited consistent force-extension characteristics (see section "Stretching of microtubule dumbbells"), so the results from ultrafast force-clamp (UFFC) experiments with differently prepared beads were combined to enhance statistical robustness.

Immobilization of pedestals and their coating with GFP-tagged proteins

Experimental chambers were assembled using silanized coverslips *(69)*, ethanol cleaned micro-slides and the double sticky tape spacers, as in *(35)*. Immobilization of 1.87 µm streptavidin-coated polystyrene beads (Spherotech, cat. # SVP-15-5) was carried out using non-specific adsorption or partial-melting methods *(35)*. Stability of the pedestals immobilized via adsorption was examined by measuring standard deviation (SD) of their thermal vibrations, and only pedestals exhibiting SD < 5 nm were used. Pedestal beads were coated with the GFP-tagged proteins using SNAP-mediated covalent binding or biotinylated anti-GFP antibodies (Abcam, cat. # ab6658) *(35)*. Briefly, for SNAP-mediated covalent binding, SNAP-GBP protein was mixed with biotinylated benzylguanine (BG) (New England Biolabs, cat. # S9110S)) for 30 min at 37 °C in ratio 25 :1 by using either 5,000 nM SNAP-GBP and 200 nM BG or 2.5 nM SNAP-GBP and

0.1 nM BG. The mixtures were incubated with coverslip-adsorbed pedestals for 30 min at room temperature. Chambers were blocked with 1% Pluronic F-127 and 1 mM biotinylated PEG, and a GFP-labeled protein diluted to 2-10 nM in phosphate buffered saline (140 mM NaCl, 2.7 mM KCl, 10.1 mM $Na_2HPO_4$, and 1.8 mM $KH_2PO_4$, pH 7.2) supplemented with 2 mg/mL bovine serum albumin (BSA) and 2 mM DTT was added for 30 min at room temperature. Pedestals immobilized via partial melting were coated with biotinylated anti-GFP antibodies (0.3 nM) and blocked with 1 mM biotinylated PEG (Quanta BioDesign, cat. # 721431-18-1) and 22.5 μM biotinylated BSA (Sigma, cat. # A8549-10MG). GFP-labeled proteins were incubated as above, but higher protein concentrations were needed to achieve similarly low percent of interacting pedestals (e.g. Ndc80-GFPc was used at 150 nM). Heating during the pedestal immobilization process appears to enhance the non-specific binding of proteins to the surface of the pedestal beads. The UFFC results from two methods of pedestal immobilization were consistent, so the data were combined to improve statistical power.

Assembly of microtubule dumbbells

All experiments were performed at 32 °C. Taxol-stabilized microtubules labeled with rhodamine and DIG were diluted 400-fold in warm Motility buffer containing Mg-BRB80 (80 mM PIPES pH 6.9, 1 mM EGTA, 4 mM $MgCl_2$), 4 mg/ml BSA (Sigma-Aldrich, cat. # A7638), 2 mM DTT, 15 μM taxol, 6 mg/ml glucose (Sigma-Aldrich, cat. # G8270), 20 μg/ml catalase (Sigma-Aldrich, cat. # C40), 0.1 mg/ml glucose oxidase (Sigma-Aldrich, cat. # G2133) and 0.5% β-mercaptoethanol. Microtubules were flowed into a chamber containing immobilized pedestals coated with Ndc80c or other protein. Dumbbell beads (4 μl) were carefully added to one side of the flow chamber to minimize excessive mixing of microtubules and floating beads (fig. S1A). The chamber was sealed with silicone rubber (Smooth-on, cat. # 10006546) and placed on a microscope stage prewarmed to 32 °C. Two beads were captured using two trapping beams and the microscope stage was moved to bring these beads to the area free from other floating beads. The captured beads were visualized briefly in the rhodamine fluorescence channel to verify absence of any bound microtubules. Calibrations of the stiffness of two traps and their corresponding quadrant photodetectors were carried out 2 μm from the coverslip surface. The stage was then adjusted to bring one of the trapped beads close to a floating microtubule (8-12 μm long) viewed via rhodamine fluorescence, facilitating the attachment of the bead to a nearby microtubule end. The microscope stage was moved repeatedly until the second trapped bead attached near the opposite end of the microtubule. Subsequently, the microtubule dumbbell was stretched by moving one of the traps along the dumbbell axis, as described in section 3.6.3 of *(35)* to achieve 2 pN pretension.

UFFC instrument description

Our microscope features differential interference contrast optical components coupled with the light emitting diode (565 nm, Thorlabs cat. # M565D2), and the epi-fluorescence light path with a high-speed shutter (Melles Griot, cat. # 04UTS201). This setup allows brightfield and epifluorescence imaging via the filter cubes (Semrock) and either a 488 nm laser (Coherent, Sapphire 488-20/460-CDRH) or a Zeiss mercury lamp for illumination. The optical trap apparatus and our ultrafast force-clamp spectroscopy setup were as in *(35)*. Briefly, a beam of the 1064 nm laser (IPG Photonics, YLR-10-1064-LP) was passed through an acousto-optic deflector (IntraAction Corp., DTD-274HA6 2-AXIS) and spilt into two separate beams via a polarizing beam-splitter cube. These beams were integrated into an upright Zeiss AxioImager.Z2 with a 100x 1.46NA oil objective, creating two optical traps. Simultaneous trap movement was accomplished by modulating AOD frequency via the field programmable gate array board-generated analog signals. The stiffnesses of optical traps typically ranged from 0.045–0.1 pN/nm (most experiments were carried out at either 45 or 90 pN/μm), and the traps had nearly identical stiffness. Position detection of trapped beads with sub-nanometer spatial precision and ~ 10 μs temporal resolution was accomplished using back focal plane detection with two independent tracking lasers of 780 nm and 830 nm (Qioptiq) paired with the custom-made QPDs. Positional coordinates (x,y,z) of the pedestal bead were monitored using a dedicated QPD at a sampling rate of 1 kHz. Nanometer-precise sample positioning was achieved using a three-axes piezo stage (Physik Instrumente P-561.3DD), which was mounted on a motorized x,y stage (ASI, PZU-4004) that was used for coarse position adjustment. Stage drifts were minimized by employing a feedback system involving a third tracking beam from a 905 nm laser (50 mW, World Star Tech, cat. # TECIR 905) focused on the pedestal bead. Calibration of the optical traps and position detectors was carried out as in *(35, 70)*. Monitoring of the pedestal bead was performed using a data acquisition card (National Instruments, cat. # PCI-6070E), and dumbbell bead positions were collected using a field programmable gate array board (National Instruments, cat. # PCIe-7842R). Description of programs to run this instrument and the codes, as well as example files with experimental raw data and data analysis, are provided in the Data source (see "UFFC software" sheet in Data source.xlsx).

Stretching of the microtubule dumbbells

Microtubule-containing dumbbell has significant elasticity, which must be taken into account for accurate data interpretation. To determine elastic properties of microtubule dumbbell, a suspended dumbbell was stretched by fixing the position of one trap and moving the second trap in 75 nm increments, up to a total distance of 450 nm (fig. S11A,B). The motion was then reversed until the tension was fully released, and the stretch and release cycle was repeated once more while recording bead coordinates relative to the original beads' positions in the unstretched microtubule dumbbell. The tension force was calculated as the product of the bead displacement in the stationary trap and its stiffness. The extension of the microtubule dumbbell was determined by subtracting the coordinate of the bead in the moving trap from that of the bead in the stationary trap (fig. S11B). Occasionally, some dumbbells exhibited an abrupt drop in force accompanied by increased extension, indicating unstable microtubule-dumbbell links; these events were excluded from further analysis. The force-extension data points for individual dumbbells were smoothed with a moving average of 10 points (fig. S11C). The incremental difference for each point was computed, and the points with differences deviating from the mean difference value by < 1% of differences' SD were retained to generate the force-extension data set. To obtain average force-extension dependency, these data points were smoothed using the moving average of 10 points and then 5 points, and every fifth point was retained. For each extension, the force-extension points were clustered using a mean shift algorithm with the bandwidth parameter 8, and the average of each cluster was calculated. This experimental data was used to generate a quantitative force-extension dependency, which was subsequently employed to determine model parameters (see Supplementary Text, Theoretical Modeling Part 1, section 'Dumbbell force-extension relationship')

UFFC assay with Ndc80 proteins

In a typical experiment, the dumbbell was oscillated for 30 s at a constant force using the synchronous movement of two traps. The direction of oscillation was along the axis of the prestretched microtubule dumbbell, which aligned with the y-axis of the QPD. The direction of motion was reversed when the leading bead displacement exceeded the preset limit; ± 100 nm or ± 175 nm were used without affecting the dumbbell velocity (fig. S1E). Dumbbell beads' coordinates were collected every 15 µs and force acting on each dumbbell bead was updated every 30 µs; the estimated response time of our system was ~ 60 µs *(35)*. With increasing force, the velocity of the dumbbell increased linearly at 32.4 ± 0.5 µm/(s·pN) *(35)*.

Control experiments included dumbbell oscillations in Motility buffer (free dumbbell oscillations) or near the pedestal coated with GFP. To conduct ultrafast force-clamp assay with protein-coated pedestals, an immobilized pedestal was selected randomly and its stability was confirmed by measuring the standard deviation of its thermal vibrations *(35)*. The stretched microtubule dumbbell was positioned near the immobilized pedestal, and the piezo stage was adjusted vertically to bring microtubule in contact with the pedestal. Stage stabilization program was initiated using the pedestal bead *(35)*. Force-clamp was executed at $F = 4$ pN clamp force using the "leading trap feedback" regime, with some experiments employing "same trap feedback" regime to confirm that the outcome did not depend on the exact method of force clamping *(35)*. Changes in the dumbbell beads' coordinates were monitored in real time using a display. If the recording exhibited regular bead oscillations as in control measurements, the pedestal was scored as "not interacting" and another randomly-selected pedestal in this chamber was examined. If the recording exhibited deviations from the regular pattern, the clamp force $F$ was changed to 8, 12, 16, 20 pN and sometimes to 2, 3 and 6 pN to carry out measurements for 30 s at each clamped force. The output file recorded the $y$ coordinates of both dumbbell beads for each force value. We routinely worked with one chamber for up to 2 hrs, collecting data for 1–3 microtubule dumbbells and 1–15 pedestals. Additionally, GFP brightness was measured for at least 30 pedestals in each chamber to monitor levels of GFP-tagged protein on the pedestals.

Experiments with motor-coated pedestals to determine microtubule polarity

To determine polarity of microtubule in individual dumbbells, the microtubule motor proteins kinesin and dynein were employed. First, a chamber was prepared with 1.87 µm pedestal beads immobilized via partial melting. Then, kinesin or dynein were conjugated to streptavidin-coated polystyrene beads of 0.54 µm diameter (Spherotech, cat. # SPV-05-10) in an Eppendorf tube to avoid contamination of the Ndc80c-coated pedestals. Human kinesin-1 K560-GFP protein with the C-terminal 6-His tag was attached via biotinylated anti-His antibody following the protocol in *(70)*. For dynein, streptavidin-coated beads were first incubated with 0.2 µM biotinylated goat anti-mouse antibodies (Jackson Immuno Research, cat. # 115-065-003), then with 0.34 µM mouse antibodies to the dynein intermediate chain (EMD Millipore, cat. # MAB1618). The unbound streptavidin on the beads was blocked with biotinylated PEG, and the beads were incubated with dynein purified from cow brain (kindly provided by E. Holzbaur). Motor-coated beads were flowed into a chamber already containing immobilized 1.87 µm pedestal beads. Motor-coated beads were allowed to adhere non-specifically to the coverslip, subsequently serving as the motor-coated pedestals. Bonsai Ndc80-GFP was added to coat the large pedestal beads, as described in section "Immobilization of pedestals and their coating with GFP-tagged proteins". A microtubule dumbbell was assembled, and the ultrafast force-clamp assay was carried out using one of the large pedestal beads in Motility buffer supplemented with 0.8 mM Mg-ATP. The same microtubule dumbbell was then brought in contact with a smaller (motor-coated) pedestal for 30 s. The density of motor coating was sufficiently high to readily detect pulling on the suspended dumbbell by the bead-coated motors.

Microtubule decoration by Ndc80c Bonsai proteins

Rhodamine-labeled, taxol-stabilized microtubules were prepared and immobilized on silanized coverslips via anti-tubulin antibodies (Biolegend, cat. #801213), as described in *(33)*. A GFP-tagged Ndc80c protein was diluted to 100 nM in Motility buffer supplemented with 0.5 mg/ml casein (Sigma-Aldrich, cat. #C5890), 2 mM DTT, and 0.5% β-mercaptoethanol. This solution was continuously perfused into the chamber at a rate of 15 µL/min during image acquisition to improve the accuracy of measurements. Several minutes after the start of perfusion, 10 images of a single field of view were captured using the mCherry and GFP channels. Each set of images was averaged, and using the averaged images, 10-pixel-wide rectangular regions were drawn around the microtubules, excluding their tips, based on the mCherry image. These regions were then transferred to the corresponding image captured in the GFP channel. Microtubule brightness was calculated as the average fluorescence intensity within each region minus the average background intensity, which was determined using a region of the same size located near each microtubule.

Molecular Docking

A short segment of the microtubule wall, consisting of two protofilaments, each containing three head-to-tail tubulin dimers with flexible tails, was prepared and equilibrated as described in *(33)*. The structural model of the Hec1 and Nuf2 CHDs of the Ndc80c was based on PDB ID 2VE7 *(25)* following the approach outlined in *(33)*. ClusPro Dock was used to dock the Ndc80 structure onto the microtubule wall fragment *(71)*. The top Ndc80–microtubule conformations (from 1,500) were identified using electrostatic-favored criteria and binding energy calculations performed with Delphi and AMBER99SB force field. The highest-scoring conformation showed Ndc80c binding to tubulins via both CHDs, involving residues K33, K41, and K115 of the Nuf2 CHD. The second-ranked conformation closely resembled the toe-binding configuration described in *(27)*.

**Analysis of the UFFC recordings**

Histogram distribution of instantaneous velocities

During one experimental measurement, a microtubule dumbbell was oscillated near one pedestal for 30 s and the data file with *y*-coordinates for both dumbbell beads was generated. To calculate the distance change between the two beads during the measurement, their coordinates were subtracted from one another. If the force-clamp operated normally, this distance, corresponding to the dumbbell length, remained constant during the entire measurement. Measurements in which this distance decreased abruptly for > 10 nm during the first 15 s of recording, indicating a loss of pretension, were excluded. If the pretension remained constant for at least 15 s from the start of measurement, the part of recording with normal pretension was used. Measurements in which the distance between beads fluctuated significantly upon Ndc80c binding, as observed with high-density coating, were also excluded.

Subsequent analysis was conducted using the coordinates of only one of the dumbbell beads. In control experiments, the bead selected for analysis was chosen randomly. In experiments near Ndc80c pedestals, the coordinates of a randomly selected bead were first visually inspected. If the bead showed no visible segments with slower-than-normal velocity, the recording was used for further analysis, representing a non-interacting pedestal. If slow velocity segments were observed, the recording of the leading bead moving toward the microtubule plus end was used, as it exhibited less thermal noise than the recording of the leading bead moving toward the minus end ("leading trap feedback" force clamp regime). In experiments using the "same trap feedback" force clamp regime, the coordinates of the clamped bead were used. Recordings with a large number of polarity-independent, short-binding events were excluded from further analysis, as such interactions were not specific to Ndc80c pedestals (Table S1).

Bead velocity was then analyzed as in *(34, 72)*. Briefly, motions in opposite directions within each recording were parsed by identifying the force turning points (fig. S4A,B). All sweeps (single segments of unidirectional motion) in the same direction were merged into one recording (fig. S4C). Velocity was calculated as the point-by-point first derivative of each merged recording and then smoothed using a Gaussian filter with a 320-point width (4.8 ms) window to generate histograms of instantaneous dumbbell velocities (fig. S4D,E,F). In control experiments with microtubule dumbbells positioned away from pedestals or near GFP-coated pedestals, a single velocity peak was observed in each direction, referred to as "free velocity." The mean free velocity was obtained by fitting each histogram with a Gaussian function. The ratio of free velocities in different dumbbell directions was calculated, and recordings where this ratio exceeded 1.3 were excluded from further analyses (Table S1).

Fraction of interacting pedestals

For each bead recording, the velocity histogram was analyzed. A recording was scored as containing Ndc80c-microtubule interaction if the histogram showed two peaks in one direction (plus-end) and either a single peak or two closely spaced peaks in the opposite direction (minus-end). The fraction of interacting pedestals was calculated as the ratio of interacting pedestals to the total number of pedestals examined under identical conditions (Table S1). If all recordings for one pedestal, collected at different clamped force values, were excluded from the analysis, that pedestal was entirely excluded from the statistics of interacting pedestals.

Semi-automatic detection of individual segments with Ndc80c sliding

Measurements carried out near some of the Ndc80c-coated pedestals generated bead coordinate recording with the slow velocity segments and the asymmetric velocity histograms (interacting pedestals). For such measurements, the free velocity peak in the plus-end direction was fitted with the Gaussian function to find mean velocity $v$ and standard deviation $\sigma$ to calculate velocity threshold ($v - 4\sqrt{2\sigma}$) for identifying segments of Ndc80c sliding. The velocity threshold decreased with decreasing force (fig. S5A). Measurements with low free velocity $v < 135$ μm/s (corresponding to < 4 pN force) often exhibited significant overlap between the free velocity peak and the peak corresponding to Ndc80c sliding; in such cases the empirically selected threshold of 40 μm/s was used. Also, the sliding velocity of Ndc80c mutant proteins exceeded the velocity of unmodified Ndc80c Bonsai, so a different procedure for velocity threshold calculation was used. First, the free velocity peak was fit with the Gaussian function, which was subtracted from the original recording to generate the likely distribution of the Ndc80c sliding velocities. The velocity threshold was then determined as the intersection between the Gaussian fitting of the free velocity peak and the distribution of Ndc80c sliding velocities. This approach reduced the number of missed sliding events, as identified by visual inspection. Recordings in which the interacting peaks in the plus-end direction were indistinguishable from the free velocity peaks due to the high velocity of sliding were excluded from further analysis (Table S1). Recordings from pedestals with a high density of Ndc80c coating produced the histograms in which the free velocity peaks in the plus-end-direction were not visible, so these recordings were not analyzed (WT – 29 recordings, 3D Nuf2 – 20, ΔTail – 7).

Semi-automatic detection of continuous bi-directional Ndc80c sliding events

A recording from a pedestal with Ndc80c-microtubule interactions contained multiple sweeps (single segments of unidirectional motion) at slow velocity. These slow sweeps occurred in tandem, ranging from 2–3 to hundreds, depending on the pedestal and the density of Ndc80c coating (fig. S3, S9). These intervals of consecutive slow-velocity sweeps alternated with intervals showing no Ndc80c sliding (consecutive fast-velocity sweeps), suggesting that repetitive reversals in the direction of the pulling force caused the splitting of an extended interaction event, during which Ndc80c continued sliding in opposite directions without detaching. We examined whether the sequential slow-velocity sweeps during one such interval corresponded to a single sliding event by measuring the velocity of the intervening (minus-end-directed) sweeps. These sweeps exhibited detectably slower velocities than those in the minus-end direction during intervals without Ndc80c sliding in the same recording (fig. S3E). Thus, intervals with repetitive slow sweeps in the plus-end direction also contained slower-than-normal sweeps in the minus-end direction, consistent with the proposal that these intervals correspond to distinct events of continuous bi-directional Ndc80c sliding.

To detect continuous sliding events in an un-biased and quantitative manner, we calculated a point-by-point derivative of the coordinate recording for one of the dumbbell beads, selected as described in the section ‘Histogram distribution of instantaneous velocities’. The resulting velocity signal was smoothed using a Gaussian filter with 64- and 320-point width windows (fig. S6A,B). The velocity threshold, calculated as outlined in the section ‘Semi-automatic detection of individual segments with Ndc80c sliding’, was applied to both smoothed velocity signals. If at least one of the curves dropped below the threshold, the corresponding point of crossing was identified as the tentative start of a sliding segment. The tentative end of the sliding segment was defined at the next crossing with the 64-point smoothed curve, or at the crossing with the 320-point curve if it also crossed the threshold (fig. S6-S7). To determine the duration of each segment, the intersections between the two smoothed velocity curves at the beginning and end of the segment were identified. The minimum detected Ndc80c sliding segment was 2 ms, as segments < 2 ms were abundant in control recordings (fig. S5B). For interactions at small forces (< 4 pN), segments identified by the algorithm often occurred in quick succession, interrupted by only a few milliseconds. The neighboring segments with duration ≥ 2 ms were combined into a single continuous sliding event. The event duration was defined as the time from the start of the first sliding segment to the end of the last segment. A cumulative distribution of the durations of sliding events (interaction times) was generated for each recording containing at least 10 events. Each distribution was fit with a single exponential function, and only distributions with an R-squared value greater than 0.9 were used to determine the characteristic times for Ndc80c sliding in individual recordings (fig. S19A).

<u>Calculation of sliding velocity and force acting on the molecule</u>

The velocity of Ndc80c sliding during each event was calculated based on the sliding velocities of individual sweeps $v_{st}$, which was analyzed in the following manner:

$$\tilde{x}(t) = v_{st}t + B\left(1 - e^{-\frac{t}{\tau_{rel}}}\right) \tag{1}$$

where $\tau_{rel}$ is a characteristic relaxation time and $B$ is the distance traveled by the bead during the reversal-caused relaxation within microtubule dumbbell. These parameters were estimated independently for each sweep using a single restriction: $\tau_{rel} \in [0, \tau_{rel}^{max}]$, where $\tau_{rel}^{max} = 350$ μ$s$ was the experimentally-derived relaxation time. Average sliding velocity for interacting events with ≥ 5 sliding sweeps was calculated for each 30 s recording.

Force acting on the sliding Ndc80c was calculated for each experimental recording using mechanical model of the oscillating dumbbell, see Supplementary text, Theoretical modeling Part 2. These data were displayed as individual points on some of the graphs, or after binning with 1 pN step of the clamped force.

### SUPPLEMENTARY TEXT: THEORETICAL MODELLING

Friction between a sliding protein molecule and the microtubule wall can be represented using a model in which the protein translocates within a periodic energy landscape, repeatedly forming and rupturing bonds. A foundational study by Bormuth et al. employed a single-bead assay to investigate experimentally the sliding of kinesin Kip3 along microtubules under a dragging force in the presence of ADP *(41)*. Using a potential energy landscape with asymmetric wells, the authors characterized Kip3 sliding with an asymmetry parameter of $\Delta = 0.33 \pm 0.01$ nm *(41)*. In the absence of external force, single Kip3 molecules diffuse with a diffusion coefficient of $D_{\mathrm{Kip3}} = 0.0043 \pm 0.0005$ μm²/s, approximately 20-fold slower than the Ndc80c *(33)*. This slow kinetics allowed for experimental confirmation that Kip3 slides along microtubules in a hand-over-hand fashion, but the molecular mechanisms underlying its sliding asymmetry remain unknown.

Building on this work, Forth and colleagues used a similar experimental approach to examine the sliding behavior of PRC1, NuMA, and EB1 proteins *(42)*. They found that while PRC1 was insensitive to microtubule polarity, both NuMA and EB1 displayed asymmetric sliding, with reported asymmetry parameters of approximately –0.4 nm and 0.45 nm, respectively. More recently, Larson et al. reported highly asymmetric sliding velocities for purified yeast and human Ndc80 proteins *(37)*. However, the single-bead assay used in that study lacks the resolution required to measure single-molecule behavior. This distinction is crucial, as ensemble behaviors do not necessarily reflect the properties of single molecules. The frictional interface of multiple Ndc80c molecules may result from repeated cycles of individual Nd80c bond formation and rupture with the microtubule wall, producing an ensemble-dependent grip even in the absence of single-molecule sliding *(44, 73)*. Furthermore, the force-velocity relationship for Ndc80c and its corresponding asymmetry parameter were not determined, leaving a critical gap in our understanding of Ndc80 sliding behavior.

In this section, we describe our theoretical approaches for elucidating the molecular mechanisms underlying the pronounced asymmetry in the sliding velocity of single Ndc80 complex (Ndc80c) molecules under force. In Part 1, we applied a spatially asymmetric transition-state model to determine the asymmetry parameter for Ndc80c, establishing unified metrics for comparison with other sliding proteins. To gain mechanistic insight into this behavior, we adopted a Brownian dynamics approach, which describes the motion of all system components with physics-based relationships and Langevin equations. Two theoretical frameworks were developed for direct comparison with experimental data. The first framework captures the mechanical and dynamic features of the UFFC experiments, incorporating optical forces and an oscillating microtubule dumbbell (Part 2, Table S3). The second framework models interactions between a sliding molecule and its binding sites on polymerized tubulin. We first applied a traditional approach that represents these interactions as occurring within periodic potential wells, corresponding to a single binding site (Part 3, Table S4). This model adequately described the force-velocity dependencies in opposing microtubule directions; however, it required significantly deeper potential wells to account for the velocity in the plus-end direction compared to the minus-end. In Part 4, we examined the model in which the sliding molecule interacts with the microtubule via two distinct binding sites—an idea informed by the known structure of the Ndc80 complex and phenotypic analyses of Ndc80c mutants *(25, 28)*. Solutions from this model displayed an asymmetric force-velocity relationship. We verified that this solution conforms to the second law of thermodynamics and demonstrated that the unconventional plus-end-directed force-velocity relationship arises from force-dependent engagement of the second binding domain (Part 5). This analysis reveals key insights into the mechanism regulating translocation velocity, highlighting the role of force-dependent engagement of the second binding site.

### Part 1. Spatially Asymmetric Transition-State Model

We employed a one-dimensional model to characterize Ndc80c's sliding asymmetry and determine its asymmetry parameter. In this model, the Ndc80c molecule translocates via force-guided diffusion within an asymmetric periodic potential, where energy wells are tilted under a constant external force $F_{mol}$. According to Kramers' rate theory *(74)*, the force-velocity relationship is described as

$$v(F_{mol}) = k_0 L \left( \exp \frac{F_{mol}\left(\frac{L}{2} + \Delta\right)}{k_B T} - \exp \frac{-F_{mol}\left(\frac{L}{2} - \Delta\right)}{k_B T} \right) \quad (2)$$

where $k_B$ is Boltzman constant, $T$ is temperature, and the asymmetry parameter $\Delta$ represents a shift of the well's minimum from the center of the period (fig. S14A). For Ndc80c, the distance between neighboring potential wells $L$, corresponding to the binding sites on microtubule, is 4 nm *(24)*. Parameter $L$, the stepping rate of the molecule $k_0$ and diffusion coefficient of the molecule in the absence of force, $D$, are linked with the following equation:

$$D = k_0 \cdot L^2 \quad (3)$$

For the diffusion coefficient of Ndc80c along microtubules $D_{\text{Ndc80}} = 0.078\ \mu\text{m}^2 \cdot \text{s}^{-1}$ *(33)*, $k_0 \approx 4{,}875\ \text{s}^{-1}$.

We used orthogonal distance regression (ODR) to fit the eq. (2) to the averaged experimental data using SD values as weights in ODR *(75)*, bins with single clamped force values were excluded. The value of parameter $\Delta = -1.30 \pm 0.11$ nm, corresponding to ≈ 33% of the potential's period, provided the best possible fit for the sliding of Ndc80c towards both ends (fig. S14B). This fitting provided good match in the lower range of plus-end-directed force, but it underrepresented velocity toward the minus-end.

### Part 2. Theoretical description of the ultrafast force-clamp assay

The ultimate goal of the UFFC assay is to determine the force-velocity relationship for a sliding molecule. The pedestal-immobilized molecule glides along microtubule under the constant pulling force $F_{mol}$, which is exerted onto the molecule by a microtubule-dumbbell. The unidirectional motion of the dumbbell, in turn, is driven by the constant force $F_{tot}$ applied to the dumbbell via two laser traps. Since portion of the force $F_{tot}$ is used to overcome viscous drag on the moving dumbbell, $F_{mol} \neq F_{tot}$. Furthermore, full mechanical system consisting of two laser traps, two dumbbell beads attached to the microtubule and the pedestal-immobilized sliding molecule is not rigid, so all compliant elements must be taken into account for accurate calculation of $F_{mol}$. In this section, we derive equations to estimate force $F_{mol}$ acting on a sliding molecule using a mechanical model of all system components corresponding to the UFFC assay.

Brief description of system components

The model considers forces and motion along a single axis aligned with the microtubule. The microtubule, modeled as a rigid cylinder, two rigid beads, a sliding molecule, and two optical traps, all undergo linear motion along this axis in a viscous medium (fig. S12A). Dragging forces act in proportion to each object's velocity; the Stokes flow approximation is used due to the low Reynolds number (Re ≪ 1). The dumbbell beads are attached laterally to the microtubule wall, creating a torque from the stretching force on each bead, which causes microtubule bowing. The resulting increase in distance between the bead centers, referred to as "end compliance" *(76, 77)* or "linkage stiffness" *(77, 78)*, is modeled with two non-linear springs connecting the ends of the rigid microtubule to the beads ('microtubule-end springs', fig. S12B). Trapping forces acting on the dumbbell beads are modeled with Hookean springs, with one end of each spring attached to the bead surface and the other end located at the center of the corresponding trap. The protein molecule is modeled as a material point anchored by a piecewise linear spring at the system's origin.

Dumbbell force-extension relationship

Force-extension relationship of the microtubule-end springs $F_{MT}(\Delta X_{dmbl})$ was derived based on the experimental force-extension relationships for the dumbbell, $F_{dmbl}(\Delta X_{dmbl})$, see section 'Stretching of the microtubule dumbbells'. The sum of the force-induced extensions of the microtubule-end springs $\Delta X_L$ and $\Delta X_R$ is responsible for the overall microtubule dumbbell extension $\Delta X_{dmbl}$:

$$\Delta X_{dmbl} = \Delta X_L + \Delta X_R \tag{4}$$

where subscripts $L$ and $R$ correspond to the left and right dumbbell ends.

To describe $F_{dmbl}(\Delta X_{dmbl})$ quantitatively, this experimental dependency was approximated with the piecewise function containing three segments separated by points $b_{dmbl}^{weak}$ и $b_{dmbl}^{strong}$ (fig. S11D, black line). During the initial segment, the dumbbell stretches with stiffness coefficient $k_{dmbl}^{weak}$:

$$F_{dmbl}(\Delta X_{dmbl}) = k_{dmbl}^{weak} \Delta X_{dmbl}, \qquad 0 < \Delta X_{dmbl} < b_{dmbl}^{weak} \tag{5}$$

Subsequently, the stiffness of the dumbbell increases. This second segment is best described by the parabolic function with parameters $A_{dmbl}, B_{dmbl}, C_{dmbl}$:

$$F_{dmbl}(\Delta X_{dmbl}) = A_{dmbl}(\Delta X_{dmbl})^2 + B_{dmbl}\Delta X_{dmbl} + C_{dmbl}$$
$$b_{dmbl}^{weak} \le \Delta X_{dmbl} \le b_{dmbl}^{strong} \tag{6}$$

During the last segment, the stiffness of the dumbbell is higher and remains constant, $k_{dmbl}^{strong}$:

$$F_{dmbl}(\Delta X_{dmbl}) = k_{dmbl}^{strong} \Delta X + d_{dmbl}^{strong}, \qquad \Delta X_{dmbl} > b_{dmbl}^{strong} \tag{7}$$

The best fit values of these parameters are listed in Table S. We also assume that microtubule dumbbell does not resist the compression. Thus, dumbbell stiffness is 0 for $\Delta X_{dmbl} \le 0$.

The force-extension dependency $F_{MT}(z)$, where $z$ represents extension of the microtubule-end springs, was calculated as

$$F_{MT}(z) = F_{dmbl}(2z) \tag{8}$$

Friction coefficient of the dumbbell bead

Because dumbbell is oscillated close to the surface of the coverslip, we calculated friction coefficients of the dumbbell beads using approach in *(79)*:

$$\gamma_B = 6\pi\eta r\left(1 + \frac{9}{16}\frac{r}{h}\right), \qquad h = 2R + l_{MAP} \tag{9}$$

Here, $\gamma_B$ is the friction coefficient for any of the two dumbbell beads, and $h$ is the distance from the microtubule axis to coverslip surface. Values of parameters were taken from the experiment: dumbbell bead radius $r$ = 260 nm is the mean of the radii of 270 nm and 255 nm for streptavidin-coated and carboxylated beads, which were used to prepare dumbbells; radius of the pedestal bead $R$ = 935 nm; medium viscosity $\eta = 10^{-3}$ Pa·s. Distance between the pedestal-bound end of the molecule and its microtubule binding site, $l_{MAP}$, was 50 nm, roughly corresponding to the length of Bonsai Ndc80c and its connecting proteins on the surface of the pedestal bead.

Thus, $h$ = 1,920 nm, resulting in $\gamma_B \approx 5.2 \cdot 10^{-3}$ pN · s/μm.

Friction coefficient of the dumbbell microtubule

Friction coefficient of the microtubule $\gamma_{MT}$ in a moving microtubule dumbbell was calculated based on the second Newton's law for each experimental recording. First, the extensions $\widetilde{\Delta X}$ of both microtubule-end springs in the pre-stretched motionless dumbbell were calculated based on the pre-stretching force $F_{pre}$ = 2 pN applied to this dumbbell. Using the force-extension relationship and $F_{MT}(\widetilde{\Delta X}) = F_{pre}$, where $F_{MT}$ is the force acting on the microtubule from the dumbbell beads, we obtain typical value of $\widetilde{\Delta X}$ = 66 nm.

Then, for each experimental recording, the velocity $v_{st}$ of the free dumbbell motion was determined by fitting a corresponding peak in the histogram of instantaneous velocities with a Gaussian function. Finally, friction coefficient of the microtubule $\gamma_{MT}$ was calculated using the system of equations that describes motions of all dumbbell components and includes an additional equation reflecting constant distance between the two traps:

$$F_2 = F_{pre} + \frac{F}{2} \tag{10}$$

$$F_1 = F_{pre} - \frac{k_1}{k_2}\frac{F}{2} + k_1\left(2\widetilde{\Delta X} - \Delta X_1 - \Delta X_2\right) \tag{11}$$

$$F_{MT}(\Delta X_1) = F_1 + \gamma_B v_{st} \tag{12}$$

$$F_{MT}(\Delta X_2) = F_2 - \gamma_B v_{st} \tag{13}$$

$$F_2 - F_1 = F_{tot} = (2\gamma_B + \gamma_{MT}) v_{st} \tag{14}$$

$$\Delta X_1 + \Delta X_2 + \frac{F_1}{k_1} + \frac{F_2}{k_2} = 2\widetilde{\Delta X} + \frac{F_{pre}}{k_1} + \frac{F_{pre}}{k_2} \tag{15}$$

Here, $F_1$ and $F_2$ are the forces acting on the dumbbell beads from the corresponding optical traps, $F_{tot}$ is the total force applied to the dumbbell, $F$ is the clamp force in this experiment, $\Delta X_1$ and $\Delta X_2$ represent the extensions of the microtubule-end springs during free dumbbell oscillation, and subscripts 1 and 2 denote the trailing and leading ends of the dumbbell, respectively. Parameters $k_1$ and $k_2$ correspond to average stiffnesses of two traps in our experiments.

The average friction coefficient $\gamma_{MT}$ for the dumbbell microtubule was similar in both directions of oscillation, as expected, yielding an estimate of $(0.017 \pm 0.004)$ pN · s/μm, mean ± SD (fig. S12C).

Calculation of force acting on the sliding Ndc80c

For each experimental recording Ndc80c sliding velocity $v$ was determined as described in section "Histogram distribution of instantaneous velocities". Dragging force acting on the sliding molecule, $F_{mol}$, was determined by solving the system of eq. (10)–(15) with $v_{st}$ replaced by $v$, and supplemented with the following equation:

$$F_{mol} = F_2 - F_1 - (2\gamma + \gamma_{MT})v \tag{16}$$

This method was used to analyze experimental results and it was also applied to the results of the simulation of Ndc80c sliding under force with the two-site model (see Part 4). The values of $F_{mol}$ calculated using the approach provided accurate estimation of the simulation results, see Part 5.

Equations of motion for the oscillating dumbbell

In the model, microtubule dumbbell was prestretched to achieve 2 pN tension along the microtubule axis, corresponding to our routine experimental conditions. Two optical traps moved in a saw-tooth pattern in a "leading trap" force-clamp regime. Specifically, position of the leading trap was updated at 33 kHz to maintain constant force $F/2$ on the leading bead, while maintaining the constant distance $\Delta_{tr} = F/2k_{tr}$ between the center of the leading bead and corresponding optical trap. Here $k_{tr}$ is the stiffness of the leading trap.

Changes in the coordinates of the dumbbell beads and microtubule were described by a system of non-linear Langevin equations, in which the inertial terms were omitted because the motion was overdamped:

$$\gamma_B \dot{x}_B^R = k_{tr}^R (x_{tr}^R - x_B^R) - F_{MT}(\Delta X_R) + \xi_B^R \tag{17}$$

$$\gamma_{MT} \dot{x}_{MT} = F_{MT}(\Delta X_R) - F_{MT}(\Delta X_L) - F_{mol} + \xi_{MT} \tag{18}$$

$$\gamma_B \dot{x}_B^L = F_{MT}(\Delta X_L) - k_{tr}^L (x_B^L - x_{tr}^L) + \xi_B^L \tag{19}$$

Here, $x_.$ is a coordinate along the microtubule axis with subscripts $B$ corresponding to a dumbbell bead. Subscripts $MT$ and $tr$ stand for microtubule and a trap. Superscripts $L$ and $R$ correspond to the left and right beads/traps; $\gamma_B$ is the friction coefficient of a dumbbell bead, $\xi_B^R, \xi_B^L, \xi_{MT}$ are random thermal forces which were modeled as Gaussian white noise; other parameters are as in Table S3.

**Part 3. Brownian modeling of Ndc80c sliding: single-site model**

General model framework

To obtain realistic description of the Ndc80c-microtubule interactions under force we used Brownian dynamics approach. The corresponding model was combined with equations for dumbbell mechanics and motions described in Part 1 'Theoretical description of the ultrafast force-clamp assay'. As in the Arrhenius-like model in Part 2, we assume that Ndc80c translocates along a linear filament represented by a one-dimensional periodic potential with a 4-nm step size. The depth of the potential well corresponds to the energy of Ndc80c-microtubule binding, which can be linked explicitly to the rate of Ndc80c force-free diffusion. For simplicity, this model omits interactions that involve the unstructured extensions of the Hec1 subunit and tubulin tails. The model also does not consider the interaction time between the Ndc80c and microtubule. This simplification means that the Ndc80c molecule diffuses without dissociation.

Description of the Ndc80c and its microtubule binding site

Ndc80c was represented by a dimensionless point fully described by a single coordinate $x_{mol}$ along one-dimension microtubule filament (fig. S13). System's origin on the filament corresponded to the molecule's attachment site on pedestal surface (not modeled explicitly). The Ndc80c point, thereafter referred to as the Hec1 point as the only microtubule-binging site in Ndc80c, translocated along filament by hopping without detachment between periodic binding sites modeled with potential wells $U_0(y)$ along microtubule axis (fig. S13A). Position of the Hec1 point within a potential well, $y$, was calculated as

$$y = \left\{\frac{x_{mol} - x_{MT}}{L}\right\} \tag{20}$$

where curly brackets denote a fractional part of the ratio, $x_{MT}$ is the coordinate of the microtubule center (midway between dumbbell beads) relative to system's origin. Each well was described using a symmetric Gaussian function:

$$U_0(y) = -G\, e^{-\frac{(L-2y)^2}{8\sigma^2}} \tag{21}$$

Here, $G > 0$ denotes the depth of the potential well corresponding to the maximal binding energy between Hec1 and its binding site. Parameter $\sigma$ defines the width of the potential well; we chose $\sigma = 0.25$ nm to represent a localized binding site on the microtubule surface, as in *(33)*. Because in this model Ndc80c binding to microtubule only via the Hec1 point, this energy also equals the maximal binding energy between Ndc80c and microtubule. Other parameters are defined in Table S4.

Force acting on Ndc80c, $F_{mol}$, was modeled as

$$F_{mol}(x_{mol}, x_{MT}) = -\frac{\partial U_0}{\partial y}(y) \tag{22}$$

Translocation of the Ndc80c under this dragging force was described by a non-linear overdamped Langevin equation:

$$\gamma_{mol}\dot{x}_{mol} = F_{mol} - F_{mol}^{elastic}(x_{mol}) + \xi_{mol} \tag{23}$$

where $\gamma_{mol}$ is friction coefficient of the Ndc80c, $F_{mol}^{elastic}$ is the elastic force of Ndc80c stretching, $\xi_{mol}$ is random thermal force acting on Ndc80c. $F_{mol}^{elastic}(x_{mol})$ was modeled as a piecewise-linear force-extension relationship which comprises the low $k_{mol}^{weak}$ and high $k_{mol}^{strong}$ stiffness regions and a change point at $x_{mol} = b_{mol}$. This widely used description of molecular stiffness reflects both the nonlinear and strain-dependent elastic properties of elongated proteins *(80, 81)*. Values of model parameters are listed Tables S3 and S4.

Numerical calculations

The system of equations (17), (18), (19) and (23) was solved numerically with explicit Euler–Maruyama method (Higham. 2001) using time step $t_{step} = 10^{-12}$ s (Table S5). The output values of all coordinates were obtained as an average of the $t_{avg} = 15$ µs to mimic the realistic data acquisition protocol during the UFFC experiments. Total simulation time $t_{total} = 75\, ms$ was empirically chosen to obtain at least 2 sweeps of dumbbell moving under the smallest force-clamp values of 2 pN. This numerical method was implemented using the custom-written programs (see Data Source file, Simulator). The core back-end computational module was written in C++. The supporting task generator and utilities, as well as the scripts for analysis of the results, were implemented in *Python 3* (see Data Source file, Analysis).

Processing of the simulation results

Sliding velocity of the molecule in simulations was calculated using the same procedure as described in section "Calculation of sliding velocity". Force $F_{mol}$ acting on the sliding Ndc80c molecule was calculated using eq. (22) and averaged over each $t_{avg} = 15$ μs logging period. To calculate the total force acting on the microtubule dumbbell through the optical traps, $F_{tot}$, the sweeps with at least 30 time points were used. The first $\lceil 1.5\ \tau_{rel}/t_{avg} \rceil$ points of each sweep were omitted to avoid the direction-reversing transitions, and only the sweeps that contained >18 time points were used for further analysis. Total force $F_{tot}$ was calculated as follows:

$$F_{tot} = k_{tr}^{L}(x_{tr}^{L} - x_{B}^{L}) + k_{tr}^{R}(x_{tr}^{R} - x_{B}^{R}) \tag{24}$$

Results of the single-site model

The above modeling framework and model parameters listed in Tables S3 and S4 were applied to predict force-velocity curves. A good match was obtained between the experimental velocity of Ndc80c translocation in the plus-end-direction using potential well depth $G \approx 10 - 11\ k_BT$ (fig. S13B). However, motion in the minus-end-direction could be described well only with a shallower well with $G \approx 7 - 8\ k_BT$. Thus, accurate descriptions of Ndc80c translocation velocities in opposite directions are achieved by using energy wells of different depths. Because different binding energies correspond to different rates of diffusion (fig. S13C), these results also imply that in the single-site model, sliding molecule has different translocation rates at zero force, and apparent contradiction with thermodynamic requirements. These findings establish unambiguously that a single-site model cannot account for strong velocity asymmetry using a single set of parameters. While force-dependent changes at the Hec1 CHD-tubulin interface cannot be excluded as the sole contributor, the results strongly suggest that Ndc80c translocation involves a more intricate mechanism than initially anticipated.

**Part 4. Two-site Brownian model of Ndc80c sliding**

The application of a single-site model, in which Ndc80c translocates within symmetric potential wells, failed to consistently describe the asymmetric velocity of translocation (Part 3). To gain mechanistic insights into the origins of this pronounced asymmetry, we developed a two-site model in which the Ndc80 complex interacts with the microtubule wall through two distinct binding sites corresponding to the globular domains of its subunits, Hec1 and Nuf2. An electron microscopy study identified the "toe" of the Hec1 subunit as the sole site for direct contact between Ndc80c and the microtubule *(24, 27)*. However, other structural, biochemical, and cell biological studies suggest that an additional microtubule-binding interface may be formed by the Nuf2 subunit *(25, 28)*. Given the prominent plus-end-directed tilt of Ndc80c on the microtubule wall *(27, 82, 83)*, we hypothesize that force acting on microtubule-bound Ndc80c induces bending that either brings the Nuf2 subunit into contact with the microtubule wall (under the plus-end-directed force) or pulls it further away (under the minus-end-directed force). In this section, we test this hypothesis using a two-site Ndc80c binding model. Unless stated otherwise, the primary model framework, approaches, and parameter values are identical to those in the single-site model described in Part 3.

Description of the Ndc80c with two microtubule-binding sites

Microtubule-binding site in the Hec1 domain was modeled in the same manner as in the model with one binding site (Part 3). The Nuf2 domain was modeled analogously as a dimensionless point located on the rigid rod at distance $d = 3.5$ nm from the Hec1 point, roughly corresponding to distance between the centers of CHDs *(25)*. The rod pivoted around the Hec1 point in the plane containing the microtubule axis and orthogonal to the pedestal surface (fig. S15A). Rotational angle $\varphi$ from 0° to 90° corresponded to the plus-end-directed tilt of the rod and values from 90° to 180° to the minus-end-directed tilt. This planar motion was restricted by a Hookean rotational spring with rotational stiffness $k_{rot}$ and the equilibrium angle $\varphi_0 = 60°$, corresponding to the orientation of the Ndc80 shaft in the microtubule-bound complex *(24)*. Pulling force from the pedestal was applied to the distal end of the rod, causing its pivoting around the microtubule-bound Hec1 point. Force towards the microtubule plus-end brought Nuf2 domain closer to the microtubule wall, whereas the minus-end-directed force increased this distance. Because mutations in the Hec1 domain strongly reduce microtubule binding affinity of the Ndc80c *(25)*, for simplicity we assume that the Nuf2 domain does not interact with microtubule if the Hec1 point is not bound to microtubule.

Description of the potential wells for Hec1 and Nuf2 points

Each domain, Hec1 and Nuf2, was assumed to interact with its dedicated binding site on a tubulin monomer. Hec1 point translocated along the microtubule filament by hopping without full detachment between the periodic binding sites, as in the single-site model (Part 3). Nuf2 point did not translocate on its own along the microtubule. Since it was attached to Hec1 via a rigid rotatable rod, it translocated together with the Hec1 point, while the rod rotated around the Hec1 point thermally or under external force. Therefore, the binding energy between the Nuf2 point and the microtubule $U_{rot}(\varphi)$ was assumed to be defined exclusively by the angle $\varphi$, which characterizes rotation of the rod. This energy was described with a Morse-like potential, $M$:

$$U_{rot}(\varphi) = M(d \sin\varphi, G_{rot}, r_0), \tag{25}$$

where

$$M(r, G_{rot}, r_0) = -G_{rot}\left(\frac{r}{r_0}\right)^2 e^{2\left(1-\frac{r}{r_0}\right)} \tag{26}$$

Here, $r = d\sin\varphi$ is distance between Nuf2 point and microtubule, $r_0$ is the parameter characterizing the width of potential well for Nuf2 point. The value of $r_0 = 0.25$ nm was chosen similarly to the $\sigma$ value in eq. (20), following *(33)*. Parameter $G_{rot} > 0$ is the depth of the potential well (fig. S15B). Angle $\varphi$ varies between 0° and 90°. For $\varphi > 90°$, the Nuf2-microtubule binding was assumed to be negligible, i.e. $U_{rot}(\varphi) = 0$ for $\varphi > 90°$.

The torque caused by binding of the Nuf2 point to its binding site on the microtubule was

$$T_{rot}(\varphi) = -\frac{\partial U_{\mathrm{rot}}}{\partial\varphi}(\varphi) \tag{27}$$

Periodic binding sites for Hec1 point translocation with the connected Nuf2 point were modeled with energy potential $U_0(y)$ (eq. (20)), with the depth of the potential well $G$ corresponding to the maximal combined binding energy of the Hec1 site $G_{main}$ and the rotational binding energy of the Nuf2 domain $U_{rot}(\varphi)$:

$$G = G_{main} + U_{rot}(\varphi) \tag{28}$$

Equations of motion for the Ndc80c with two binding sites were the same as in the single-site model: equations (17)–(19) and (23). They were supplemented with additional equation that describes rotation of the rod:

$$\zeta\dot{\varphi} = -k_{rot}(\varphi - \varphi_0) - F_{mol}^{elastic}(x_{mol})\, l \sin\varphi + T_{rot}(\varphi) + \xi_{angle} \tag{29}$$

Here, $\zeta$ is the rotational friction coefficient, $\xi_{angle}$ represents thermal torque causing angular motion of the rod and the term $F_{mol}^{elastic}(x_{mol})\, l\sin\varphi$ represents torque from the pedestal that rotates the rod toward the microtubule. Rotational friction coefficient $\zeta$ was estimated based on *(84)* assuming that rod is represented by a cylinder of length $l$ and diameter $d_{mol} = 2$ nm:

$$\zeta = \pi\eta l^3 \left(\frac{1}{3(\log p+\delta_\perp)} + \frac{1}{\log p+\nu_\perp}\right) \quad (30)$$

where

$$p = \frac{l}{d_{mol}}, \qquad \delta_\perp = -0.662 + \frac{0.917}{p} - \frac{0.05}{p^2}, \qquad \nu_\perp = 0.839 + \frac{0.185}{p} + \frac{0.233}{p^2}$$

Numerical calculations and analysis of theoretical results were carried out as in Part 3.

**Part 5. Analysis of the Two-Site Model of Molecular Translocation**

Analysis of Force-Velocity Relationship

To investigate the behavior of the two-site model, we simulated molecular translocation under clamped forces. Representative modeling outcomes, including calculated molecular characteristics, are shown in figs. S16 and S17. To simplify the analysis of the force-velocity relationship, sliding was calculated for a constant, unidirectional force applied to the dumbbell, ranging from 2 pN to 32 pN. Additionally, to streamline this analysis, thermal noise was minimized by employing higher trap stiffness than in our typical experiments ($k_{tr}^{L,R}$ = 200 pN/µm) and the dumbbell beads attachment to microtubules was assumed to be rigid. Additional model parameters are detailed in Tables S3 and S4.

Time-dependent coordinates of the sliding molecules (fig. S16A, B) were used to determine sliding velocity and the force acting on the sliding molecule, $F_{mol}$. For consistency, the same methods used in experimental data analysis were employed for calculating sliding velocity and estimating $F_{mol}$ (see sections 'Calculation of Sliding Velocity' and 'Calculation of Force Acting on the Sliding Ndc80c'). Unlike in experiments, the force acting on the sliding molecule in simulations can be directly computed. We compared these directly computed values with those estimated using our experimental data analysis approach. Across a broad range of model parameters ($G_{main} = 4$–$7\ k_BT$, $G_{rot} = 5$–$8\ k_BT$, $k_{rot} = 10^{-3}$–$10^{-1}$ pN · µm), the average error in estimating $F_{mol}$ was less than 0.5 pN (Fig. S12D). This result supports the validity of our approach for estimating molecular force in UFFC experiments.

Force-velocity dependencies were then plotted to analyze their shapes and characteristics. The calculated force-velocity functions for sliding translocation in opposite directions exhibited notably distinct profiles. Fig. S18A illustrates this dependency for $G_{main} = 7\ k_BT$ and $G_{rot} = 5\ k_BT$, and a rotational stiffness of $k_{rot} = 0.05$ pN · µm for the rod containing the second microtubule-binding site—parameters that reduce the binding of the second site in the absence of a dragging force. The minus-end-directed force-velocity relationship displays a steep, exponential-like increase in velocity, plotted with negative values for both velocity and force. In contrast, the plus-end-directed function demonstrates a complex shape that deviates significantly from a simple exponential curve. At low dragging forces, the velocity in the plus-end direction increases rapidly, with an initial slope comparable to that of the minus-end direction (fig. S18B). This similarity validates the thermodynamic consistency of our model, confirming that velocities are direction-independent at zero external force. As the plus-end-directed dragging force increases further, the translocation velocity shows a weak dependence on the applied force. However, with higher dragging forces, velocity adopts an exponential character, albeit with a different slope than at low force corresponding to nearly constant engagement of both binding sites. These findings reveal that the two-site interaction model enables pronounced velocity asymmetry during single-molecule translocation, with distinct responses in the minus-end and plus-end directions.

Force-Dependent Engagement of the Second Microtubule-Binding Site

To investigate this unusual behavior, we computed the rotational binding energy $U_{rot}$ of the second site as a function of dragging force. During minus-end-directed motion, the rod orientation exhibits significant fluctuations, often pointing away from the microtubule plus-end (fig. S16E, S17B, Video 1). Consistently, the rotational binding energy $U_{rot}$ remains on average low (fig. S16F, S18C). At zero force, the second site's rotational binding energy is approximately $\approx 0.5\ k_B T$, arising from occasional thermal fluctuations. This indicates that microtubule binding by the second site during minus-end-directed translocation is minimal. To test this conclusion further, we computed the force-velocity relationship for translocation using a single-site model, with all parameter values identical to those in the two-site model. The energy well depth in the single-site model matched the binding energy of the first (Hec1) site in the two-site model ($7\ k_B T$). The single-site model produced a force-velocity relationship in the minus-end direction closely aligning with the predictions of the two-site model (fig. S18A). These results demonstrate that minus-end-directed sliding primarily depends on the Hec1 site, with minimal engagement of the Nuf2 site. This reduced engagement explains the fast velocity and the exponential-like character of the minus-end-directed force-velocity relationship.

During motion in the plus-end direction, the angle between the rod and the microtubule remained very small (fig. S16E). Correspondingly, binding of the second domain was pronounced (fig. S16F) and increased steadily with rising force (fig. S18C). At approximately 6 pN of plus-end-directed force, the sliding molecule reached its maximum friction-generating configuration. At this point, the sliding velocity began to increase as the second site's capacity to buffer velocity became exhausted. Within this force range, the force-velocity relationship transitioned to an exponential-like regime, and the friction coefficient started to decline (fig. S18D). Interestingly, the rotational binding energy $U_{rot}$ of the second site appeared to approach a maximum value of approximately $4.2\ k_B T$, which was lower than the expected maximum ($G_{rot} = 5\ k_B T$) (fig. S18C). This discrepancy likely arises from thermal fluctuations of the translocating molecule, which prevent the second site from maintaining consistent binding. Moreover, at higher forces, these fluctuations also impact the potential depth of the first site, suggesting that the two-site molecular translocation operates as a fluctuating potential ratchet (Reimann, 2002).

To further evaluate the force-velocity relationship for translocation in the plus-end direction, we employed a single-site model with an energy well depth representing the combined binding energies of both the Hec1 and Nuf2 sites in the two-site model ($11.2\ k_B T$). This energy, corresponding to full engagement of both sites, yielded the expected exponential-like velocity profile (fig. S18A). These results confirm that the complex shape of the plus-end-directed force-velocity dependency in the two-site model arises from the force-dependent engagement of the second site. Importantly, the exact shape of the force-velocity relationship and the range of forces for maximal friction coefficient increase depend on molecular parameters, such as the binding energies of the two sites and molecular stiffness. For the human Ndc80 complex studied experimentally in our work, the friction-modulating regime occurs at lower forces than in the model solution examined in this section. Future studies should explore whether this operational range can be tuned by Ndc80c-binding proteins or posttranslational modifications, which presents an intriguing question for further investigation.

### Supplementary Tables

**Table S1. UFFC experimental statistics.** Experiments were carried out using pedestals coated with different GFP-tagged Ndc80c Bonsai proteins. WT refers to unmodified Ndc80c Bonsai protein. MT-microtubule.

| Experimental statistics | WT | Ndc80c-3D Nuf2 | Ndc80c-Δ80 | Ndc80c-Hec1 K166D |
|---|---|---|---|---|
| number of experimental chambers | 43 | 12 | 6 | 8 |
| number of pedestals examined with MT dumbbell | 347 | 110 | 95 | 116 |
| not interacting | 255 | 80 | 84 | 88 |
| interacting | 59 | 20 | 7 | 11 |
| excluded for reasons described in section "Analysis of the UFFC recordings" | 33 | 10 | 4 | 17 |
| fraction of interacting pedestals per chamber | 0.19 | 0.20 | 0.08 | 0.11 |
| number of measurements (30s recordings) | 620 | 149 | 129 | 178 |
| number of measurements with no interaction | 351 | 84 | 86 | 93 |
| number of measurements excluded from analysis for reasons listed below | 99 | 15 | 8 | 34 |
| loss of MT dumbbell pretension | 33 | 7 | 4 | 1 |
| unusual fluctuations in MT dumbbell length | 5 | 1 | 0 | 7 |
| non-specific sticking to pedestal | 31 | 0 | 0 | 4 |
| high free velocity ratio of the dumbbell beads | 30 | 5 | 4 | 16 |
| interacting peaks are too close | 0 | 2 | 0 | 6 |

**Table S2. Brownian model of the UFFC assay: model variables.** List of all variables used in section "Theoretical modelling". MT – microtubule.

| Symbol | Description | Units |
|---|---|---|
| $F_{mol}$ | dragging force acting on the MT-bound molecule | pN |
| $F_{tot}$ | total force applied to the dumbbell via optical traps | pN |
| $F_1, F_2$ | forces exerted by the optical trap on the dumbbell beads, with subscripts 1 and 2 denoting the trailing and leading ends of the dumbbell, respectively | pN |
| $\Delta X_{dmbl}$ | MT dumbbell extension | μm |
| $k_1, k_2$ | stiffness of the trailing (1) and leading (2) traps | pN/μm |
| $\Delta X_{L,R}$ | extensions of the left (L) and right (R) microtubule-end springs | μm |
| $\widetilde{\Delta X}$ | extension of the microtubule-end spring in a pre-stretched dumbbell | μm |
| $\Delta X_{1,2}$ | extensions of the microtubule-end springs during free dumbbell oscillation, subscripts 1 and 2 denoting the trailing and leading ends of the dumbbell, respectively | μm |
| $\Delta_{tr}$ | distance between the leading trap and the leading bead during oscillation in the force clamp regime | μm |
| $x_B^{R,L}$ | coordinates of the right (R) and the left (L) beads relative to the center of the pedestal | μm |
| $x_{tr}^{R,L}$ | coordinates of the right (R) and the left (L) traps relative to the center of the pedestal | μm |
| $\tilde{x}_B^{R,L}$ | displacement of the right (R) and the left (L) beads from the centers of the corresponding traps | μm |
| $x_{MT}$ | coordinate of the MT center (midpoint) | μm |
| $x_{mol}$ | coordinate of the molecule along microtubule filament | μm |
| $\tau_{rel}$ | relaxation time of the MT dumbbell after directional change | μs |
| $v_{st}$ | stationary velocity of free dumbbell motion in experiments | μm/s |
| $v$ | velocity of molecule sliding along the MT | μm/s |
| $\xi_B^R, \xi_B^L$, $\xi_{MT}, \xi_{mol}$ | thermal forces acting on the right and left beads, the MT and the molecule | pN |
| $\xi_{angle}$ | thermal torque acting on the Ndc80c in the two-site model | pN·μm |
| $y$ | molecular coordinate within the potential well | μm |
| $\varphi$ | angle between Ndc80c rod and MT axis in the two-site model | ° |
| $r$ | distance between the second (Nuf2) binding site and microtubule | μm |

**Table S3. Brownian model of the UFFC assay: parameters and functions describing experimental configuration.**

| Symbol | Description | Unit | Value | Source |
|---|---|---|---|---|
| $\gamma_{MT}$ | MT drag coefficient | pN·s/µm | 0.017 (mean) | estimated based on experimental measurements, see section 'Friction coefficient of the dumbbell microtubule' |
| $\eta$ | medium viscosity | Pa·s | $10^{-3}$ | water viscosity |
| $\gamma_B$ | drag coefficients of the dumbbell bead | pN·s/µm | $5.2 \times 10^{-3}$ | analytically calculated, see section 'Friction coefficient of the dumbbell bead' |
| $T$ | temperature | K | 300 K | experimental parameter |
| $F_{dmbl}(z)$ | force-extension dependency of the MT dumbbell | pN | eq. (5)–(7) | derived based on experimental measurements |
| $F_{MT}(z)$ | force-extension dependency of the MT end-spring | pN | eq. (8) | derived based on experimental measurements |
| $F_{pre}$ | pre-stretch force applied to the MT dumbbell | pN | 2 | experimental condition |
| $F$ | clamp force | pN | 2–32 (varied) | experimental condition |
| $k_{tr}^{L}$ | stiffness of the left (L) optical trap | pN/µm | 42 | experimental condition |
| $k_{tr}^{R}$ | stiffness of the right (R) optical trap | pN/µm | 44 | experimental condition |
| $l_{MT}$ | length of the MT | µm | 9 | typical distance between the dumbbell beads in experiments |
| $R$ | radius of pedestal bead | nm | 935 | experimental condition |
| $r$ | radius of dumbbell bead | nm | 260 | experimental condition |
| $l_{MAP}$ | distance between the pedestal-bound end of the molecule and its MT binding site | nm | 50 | approximate length of Ndc80c Bonsai molecule and SNAP-GBP linker |
| $h$ | distance from the MT axis to the coverslip surface | nm | 1,920 | experimental parameter |

**Table S4. Brownian model of the UFFC assay: Ndc80c interaction with microtubule**.

| Symbol | Description | Unit | Value | Source |
|---|---|---|---|---|
| $D_{Ndc80}$ | Ndc80c diffusion coefficient | µm²/s | 0.078 | *(33)* |
| $\gamma_{mol}$ | drag coefficient of Ndc80c | pN·s/µm | $1.38 \cdot 10^{-4}$ | this work, analytically estimated (see section viscous drag forces on the beads and the molecule') |
| $F_{mol}^{elastic}(x_{mol})$ | force-extension characteristic of Ndc80c | – | – | see section 'Description of the Ndc80c and its microtubule binding site' |
| $\sigma$ | width of the potential well of the first site | nm | 0.25 | *(33)* |
| $L$ | periodicity of Ndc80c binding sites | nm | 4 | *(27)* |
| $G$ | total MT-Ndc80c binding energy | $k_B T$ | 0–15 (varied) | this work |
| $U_0(y)$ | periodic potential energy function | $k_B T$ | eq. (20) | this work |
| $l$ | length of Ndc80c | nm | 17 | Ndc80 Bonsai *(25)* |
| $\varphi_0$ | tilt angle of the MT-bound Ndc80c | ° | 60 | *(24)* |
| $d$ | distance between the Hec1 and Nuf2 points along the rod | nm | 3.5 | see section 'Description of the Ndc80c with two microtubule-binding sites' |
| $r_0$ | width of potential well of the second site | nm | 0.25 | this work |
| $\zeta$ | rotational drag coefficient of Ndc80c | pN·µm·s | $2 \cdot 10^{-7}$ | this work, eq. (30) |
| $G_{main}$ | depth of potential well of the first site | $k_B T$ | 0–9 (varied) | this work |
| $G_{rot}$ | depth of potential well of the second site | $k_B T$ | 0–8 (varied) | this work |
| $M(r, G_{rot}, r_0)$ | Morse-like potential of interaction between the second site and MT | $k_B T$ | eq. (26) | this work |
| $U_{rot}(\varphi)$ | potential energy well for rotational motion | $k_B T$ | eq. (25) | this work |
| $T_{rot}(\varphi)$ | torque between the second site and MT | pN·µm | eq. (27) | this work |
| $k_{rot}$ | rotational stiffness of Ndc80c | pN·µm | $10^{-5}$–$10^{-2}$ (varied) | this work |

**Table S5. Parameters of numerical simulations.**

| Parameter | Description | Unit | Value | Source |
|---|---|---|---|---|
| $t_{step}$ | Time step for numerical calculation | s | $10^{-12}$ | See section "Numerical calculations" |
| $t_{avg}$ | Averaging and logging time step | μs | 15 | Matches data acquisition time in our experiments |
| $t_{total}$ | Total simulation time for UFFC experiments | ms | 75 | See section "Numerical calculations" |

### Supplementary Video legend

**Video 1. Direction-dependent engagement of the Nuf2 CHD in the sliding Ndc80c molecule.**

This video shows results from theoretical simulations using the two-site model with the following parameters: clamp force 4 pN, $G_{main}$= 6 $k_B T$, $G_{rot}$= 6 $k_B T$, $k_{rot} = 10^{-2}$pN · μm, and a time step 1 ps. Frames represent averaged coordinates and angles over 15 μs and are played at 5 fps, corresponding to 13,300× slower playback than real time. In the video, the Hec1 CHD (blue) and Nuf2 CHD (yellow) of Ndc80c are connected to a stalk fragment, while other components of the full-length Ndc80c are omitted for clarity. A single microtubule protofilament is shown with repetitive binding sites (4 nm periodicity) represented as complementary, cognate regions specific to the two CHDs. Initially, an external force is applied in the plus-end direction, during which the Nuf2 CHD remains almost continuously bound to the microtubule, except for fast directional hops with 1–3 tubulin monomer steps. Reversing the force disengages the Nuf2 CHD, resulting in rapid sliding toward the minus end. Detailed results of this simulation are shown in fig. S17.

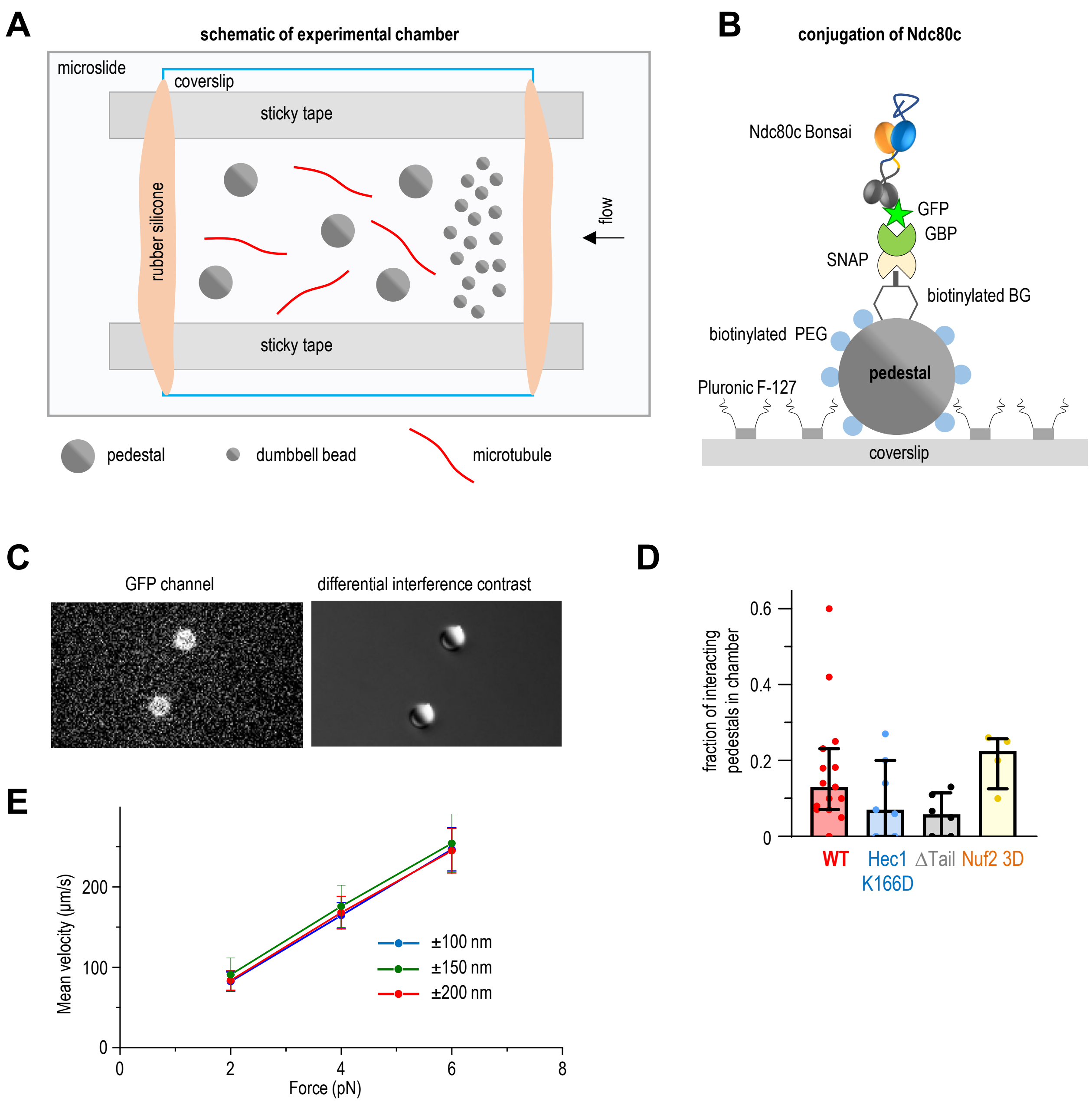


**Fig. S1. Experimental details of the UFFC assay. (A)** Schematic of the assembled microscopy chamber for the UFFC assay (not to scale). **(B)** Schematic of the Ndc80c conjugation to pedestals using SNAP-GBP linker (not to scale), see Materials and Methods for more details. **(C)** GFP-fluorescence and DIC images of the coverslip-immobilized pedestals coated with Bonsai Ndc80c-GFP. **(D)** Fraction of interacting pedestals. Each dot represents results for an individual chamber with at least 10 tested pedestals. Bars represent median values, whiskers - 25%-75% interquartile range. A two-tailed t-test showed that difference between these groups is not significant. **(E)** Mean (free) velocity of microtubule dumbbells with SD (N =3 for each data point) tested in the absence of pedestals.

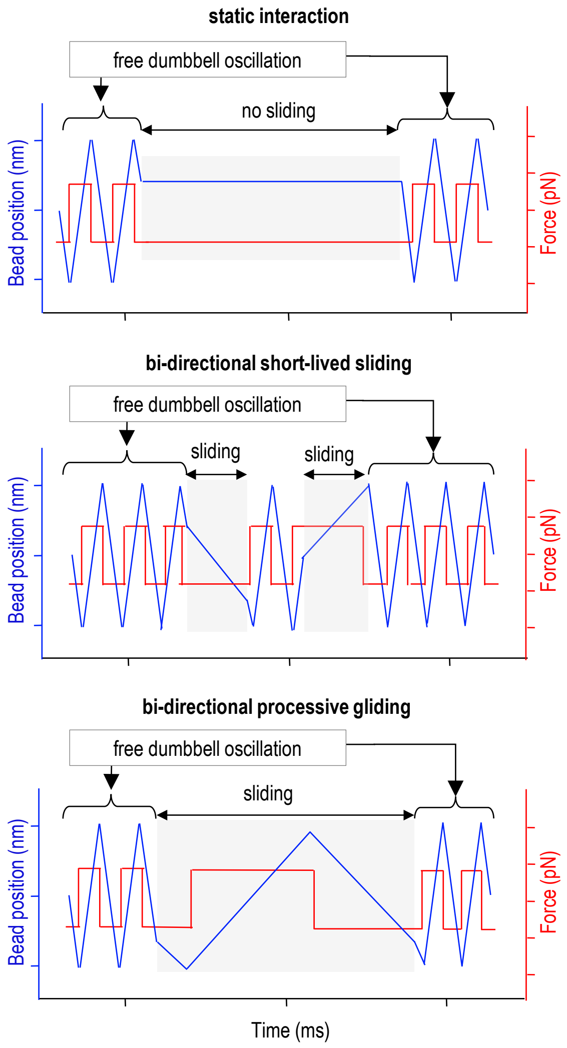


**Fig. S2. Schematized recordings illustrating possible outcomes of the UFFC assay.** The position of a dumbbell bead (blue) and applied force (red) are shown for different Ndc80c interactions with microtubules, varying in binding strength and sliding ability. Free dumbbell oscillations appear as repetitive “triangles,” while molecular interactions are highlighted in gray. Sharp bead trajectory changes reflect the lack of system compliance in these illustrative traces. In actual systems, dumbbell compliance produces smooth, non-linear transitions. The short length of the Ndc80c Bonsai protein (< 20 nm) minimizes the impact of molecular "flipping" during abrupt directional changes.

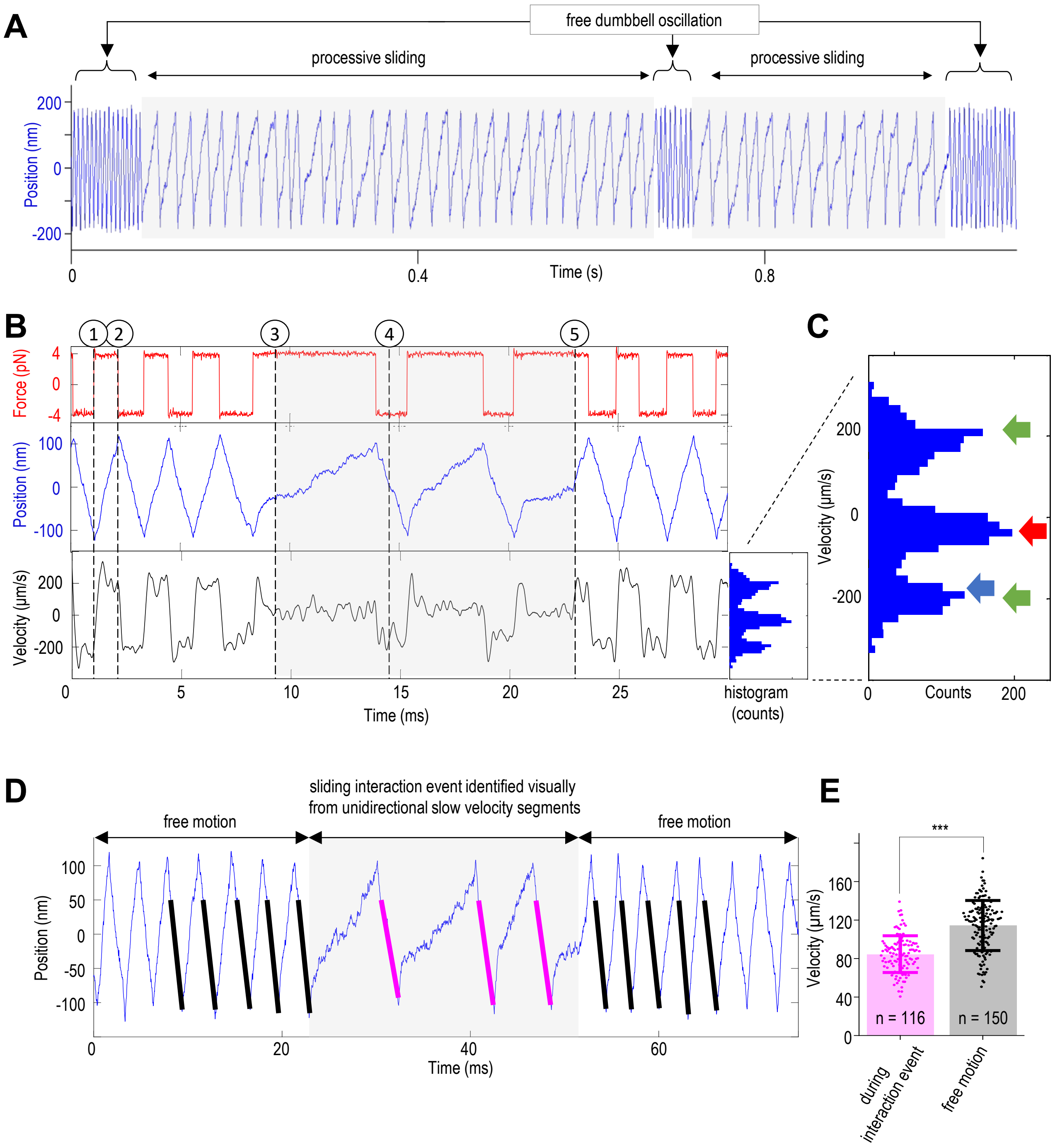


**Fig. S3. Typical recordings for the Ndc80c sliding.** Bonsai Ndc80c-GFP was investigated using the "same trap feedback" regime at 4 pN clamped force. **(A)** A fragment of the recording (1 s) showing changes in position of one of the dumbbell beads (blue) and two interaction events. **(B)** A 30 ms fragment with one interaction event additionally showing changes in force applied to the microtubule dumbbell (red), changes in velocity of this bead obtained from point-by-point differentiation and smoothing with a Gaussian filter with 320-point width window (black) and the corresponding instantaneous velocity distribution. Vertical dashed lines mark distinctive signal features: 1 – free dumbbell motion, 2 – reversal of the direction of clamped force, 3 – start of the slower velocity segments (Ndc80c sliding), 4 – Ndc80c sliding in opposite direction with faster velocity, 5 – unbinding of Ndc80c and resumption of free oscillations. **(C)** Enlarged distribution of instantaneous velocities in the example signal shown in panel B. Green arrows indicate free velocity peaks in two different directions, red arrow indicates the plus-end-directed Ndc80c sliding peak, blue arrow indicates approximate position of the minus-end-directed velocity peak, which is positioned sloe to the free velocity peak. **(D)** Manual analysis of the continuity of Ndc80c

**Fig. S3** (continued) sliding upon oscillating force. Example segment of a bead position for Ndc80c Bonsai at 4 pN clamp force, highlighting a tandem sequence with slow individual sweeps in the plus-end direction (gray box). Linear fits are shown for individual sweeps in the minus-end direction within the tandem sequence (magenta) and for five sweeps immediately before and after the sweeps in the gray box (black lines), representing control free dumbbell motion. The initial 50 nm segments of each sweep were excluded from the fit, as they correspond to dumbbell relaxation following abrupt directional reversal. **(E)** Dots show individual sweep velocities determined from the slopes of linear fits, such as shown in panel D, for one 30 s measurement. Bars and whiskers are Mean with S.D.; unpaired t-test, ***$p < 0.001$. Reduced velocity of the fast segments alternating with the slow segments confirms that Ndc80c remains bound to microtubule as it slides in both microtubule directions.

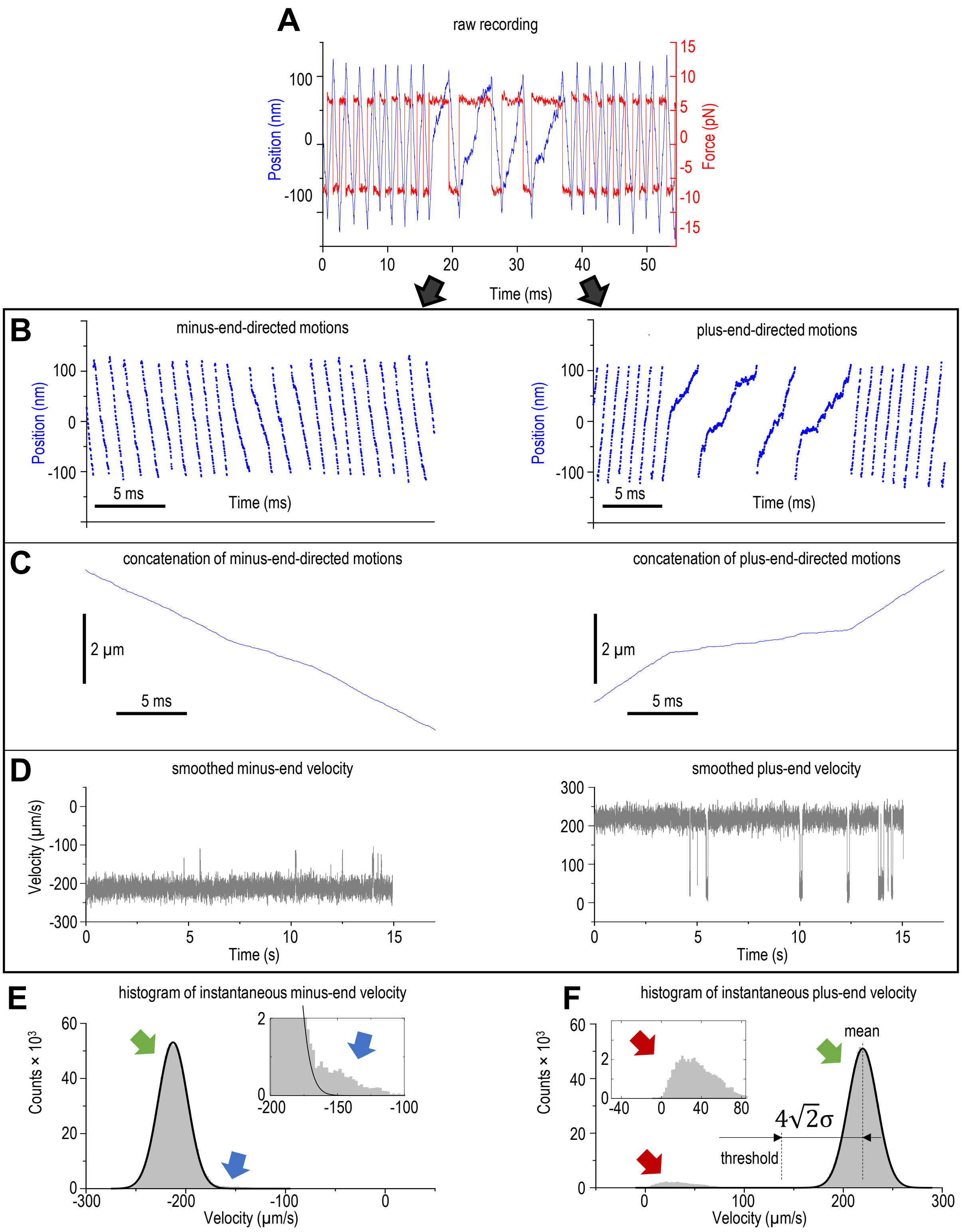


**Fig. S4. Construction of instantaneous velocity histograms. (A)** A fragment (55 ms) of the example recording for Ndc80c Bonsai at 6 pN force. Panels B-D show analysis of motions toward the microtubule minus-end (left) and plus-end (right). **(B)** Unidirectional bead coordinates for individual sweeps. **(C)** Unidirectional concatenated recordings. **(D)** Changes in velocity for the concatenated recording for the entire 30 s experimental signal. Velocity was calculated by differentiating coordinate data point-by-point and applying Gaussian smoothing with a 320-point window. **(E)** and **(F)** show instantaneous velocity histograms with enlarged insets. Green arrows point two free velocity peaks, black lines show Gaussian fitting. Red and blue arrows point to Ndc80c-dependent velocities.

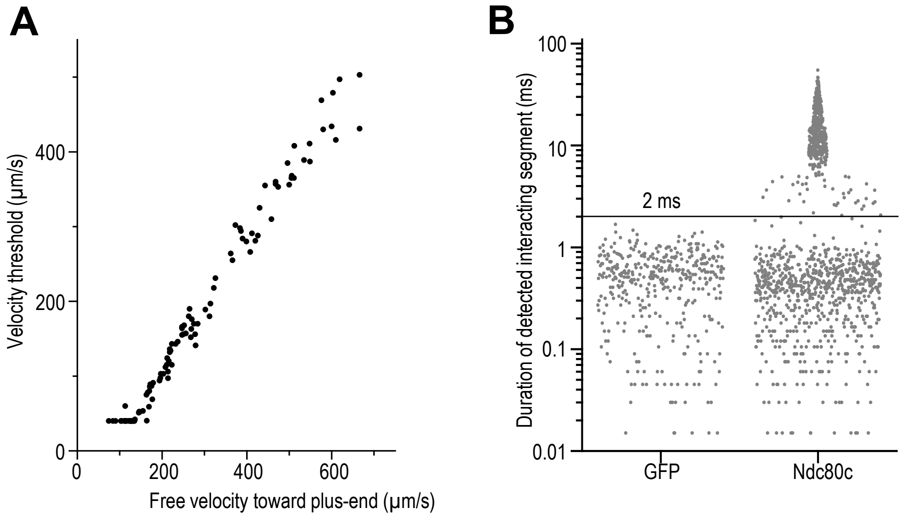


**Fig. S5. Semi-automatic detection of Ndc80c sliding segments. (A)** Velocity threshold as a function of free velocity was plotted based on data for n = 106 recordings obtained in N = 43 chambers. Each point depicts a threshold value determined for one 30 s recording. **(B)** Durations of sliding segments detected by the semi-automatic detection algorithm for pedestals coated with GFP (control, 402 segments detected in 2 recordings) and Ndc80c Bonsai (1,185 segments detected in 2 recordings) at 4 pN force. Virtually all selected segments for GFP were < 2 ms (horizontal black line), so this time was used as a cut-off for the algorithm.

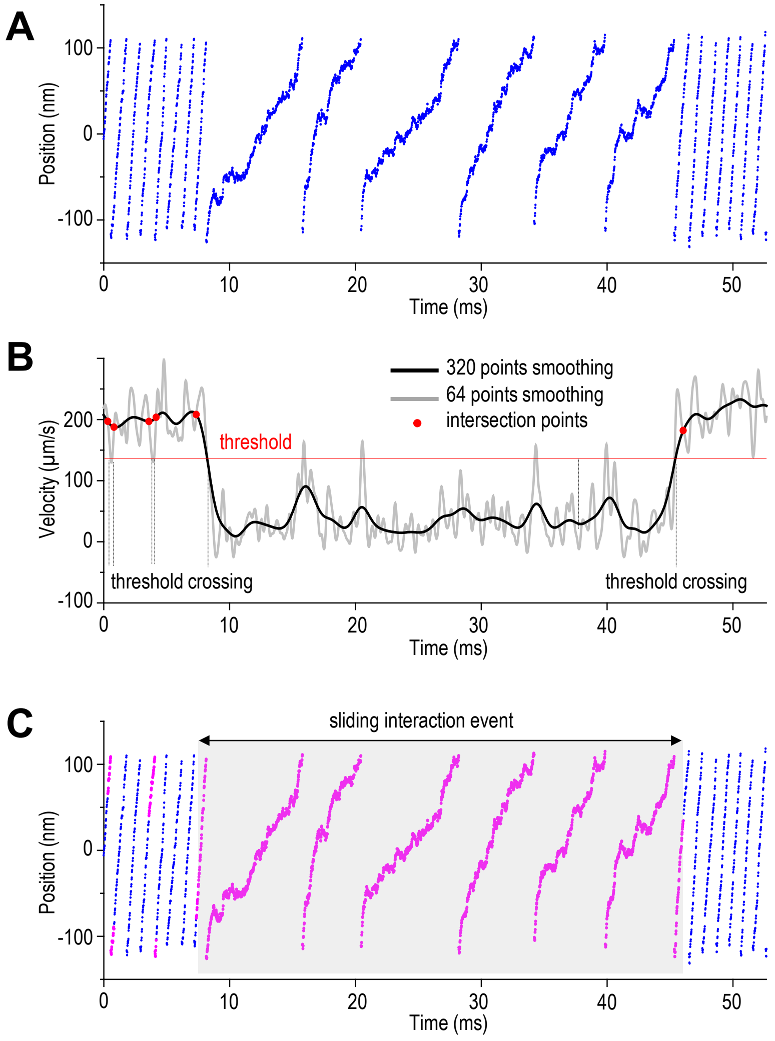


**Fig. S6. Semi-automatic detection of continuous interaction events (7 pN clamp force). (A)** A fragment of the bead position recording with the plus-end-directed tandem sweeps at slower velocity; data for Ndc80c Bonsai at 7 pN. **(B)** Instantaneous velocity for bead recording in panel A, see "Materials and Methods" section "Analysis of the UFFC recordings". Gray curve – point-by-point velocity was smoothed by applying Gaussian filter with 64-point window; black – 320-point window. Red horizontal line at 140 µm/s marks the velocity threshold calculated as described in section "Semi-automatic detection of individual segments with Ndc80c sliding". Red dots indicate velocity curves intersections that are proximal to their crossings of red line. **(C)** Same bead position recording as in panel A but showing individual sliding segments (in magenta) identified by the algorithm. Consecutive segments that lasted > 2 ms were combined in one sliding interaction event (gray box). All individual sliding segments outside the gray box were < 2 ms, so they were excluded.

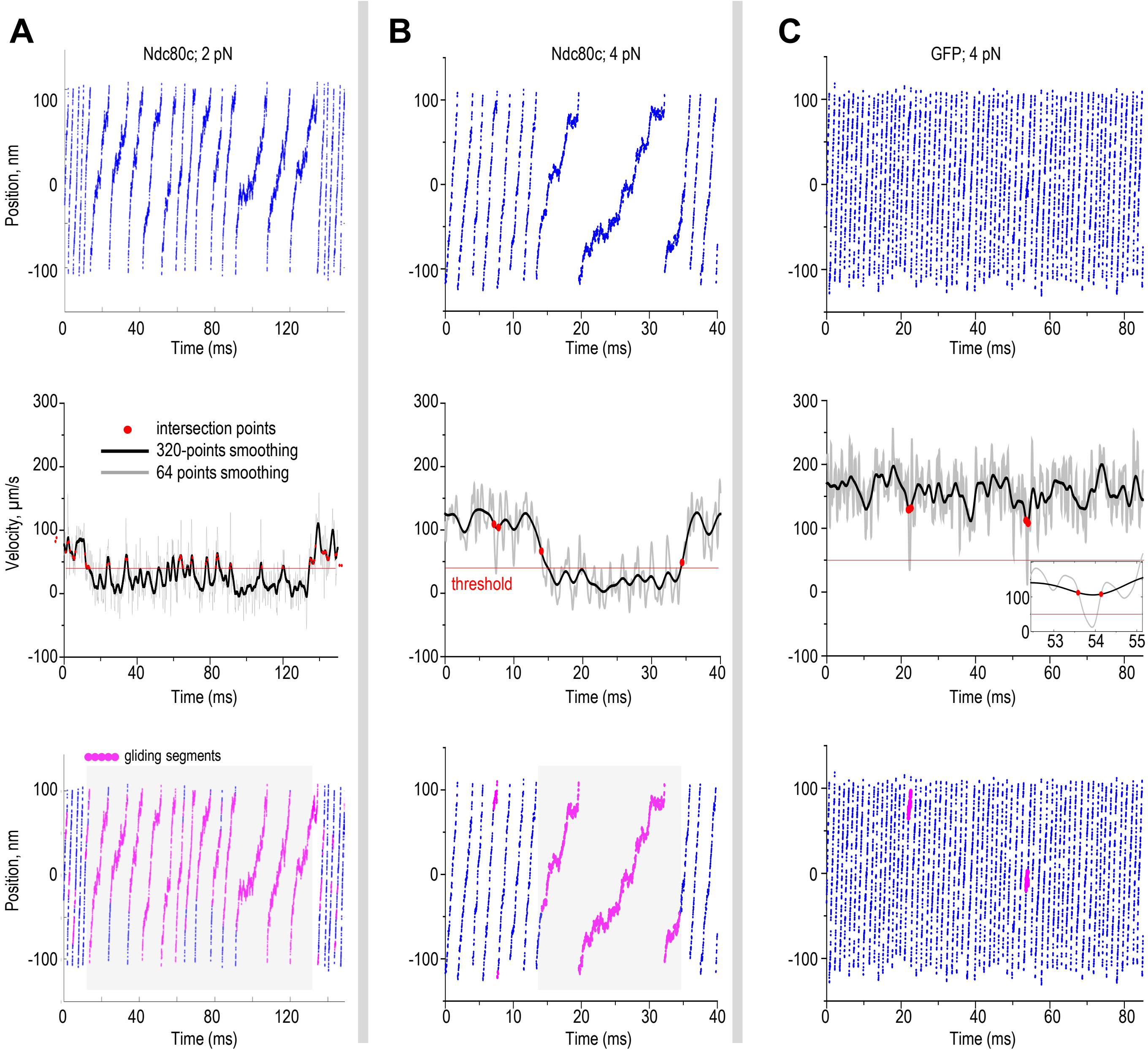


**Fig. S7. Semi-automatic detection of continuous interaction events.** Examples of interaction events recorded for Ndc80c Bonsai at 2 pN (**A**) and 4 pN (**B**) clamped force. (**C**) Example recording obtained with GFP-coated pedestals at 4 pN clamp force. See legend for fig. S6 for details.

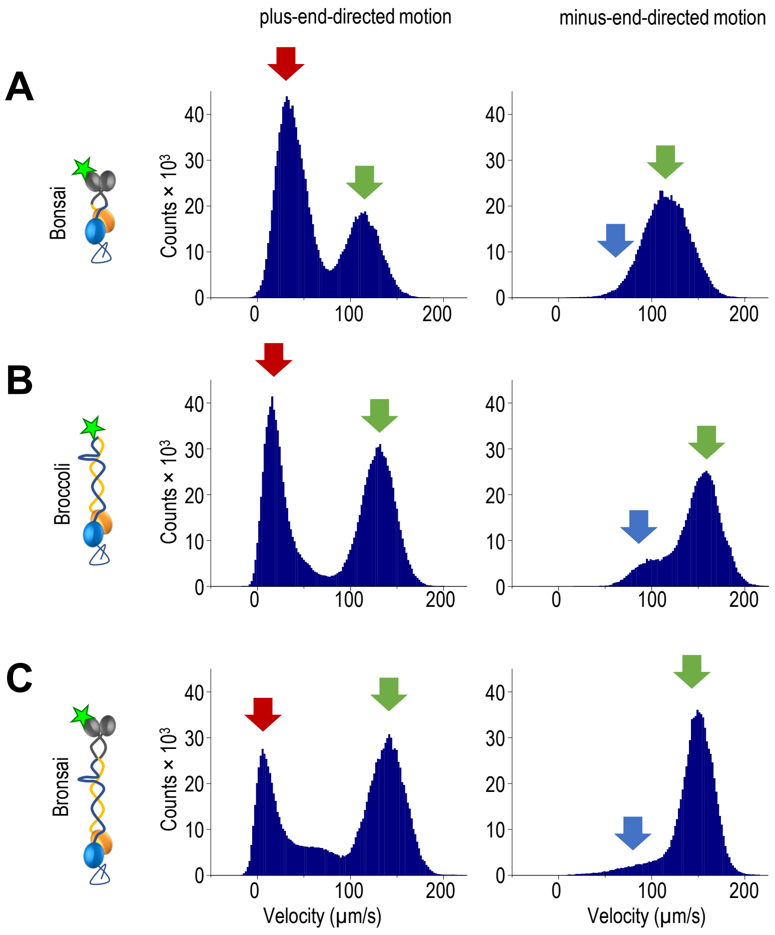


**Fig. S8. Asymmetric sliding of different Ndc80c constructs.** Different Ndc80c constructs were tested analogously in the UFFC assay and example velocity histograms were plotted. See legends to fig. S3C. Minor differences in the appearance of histograms obtained under identical conditions represent experimental variability seen with each of these proteins.

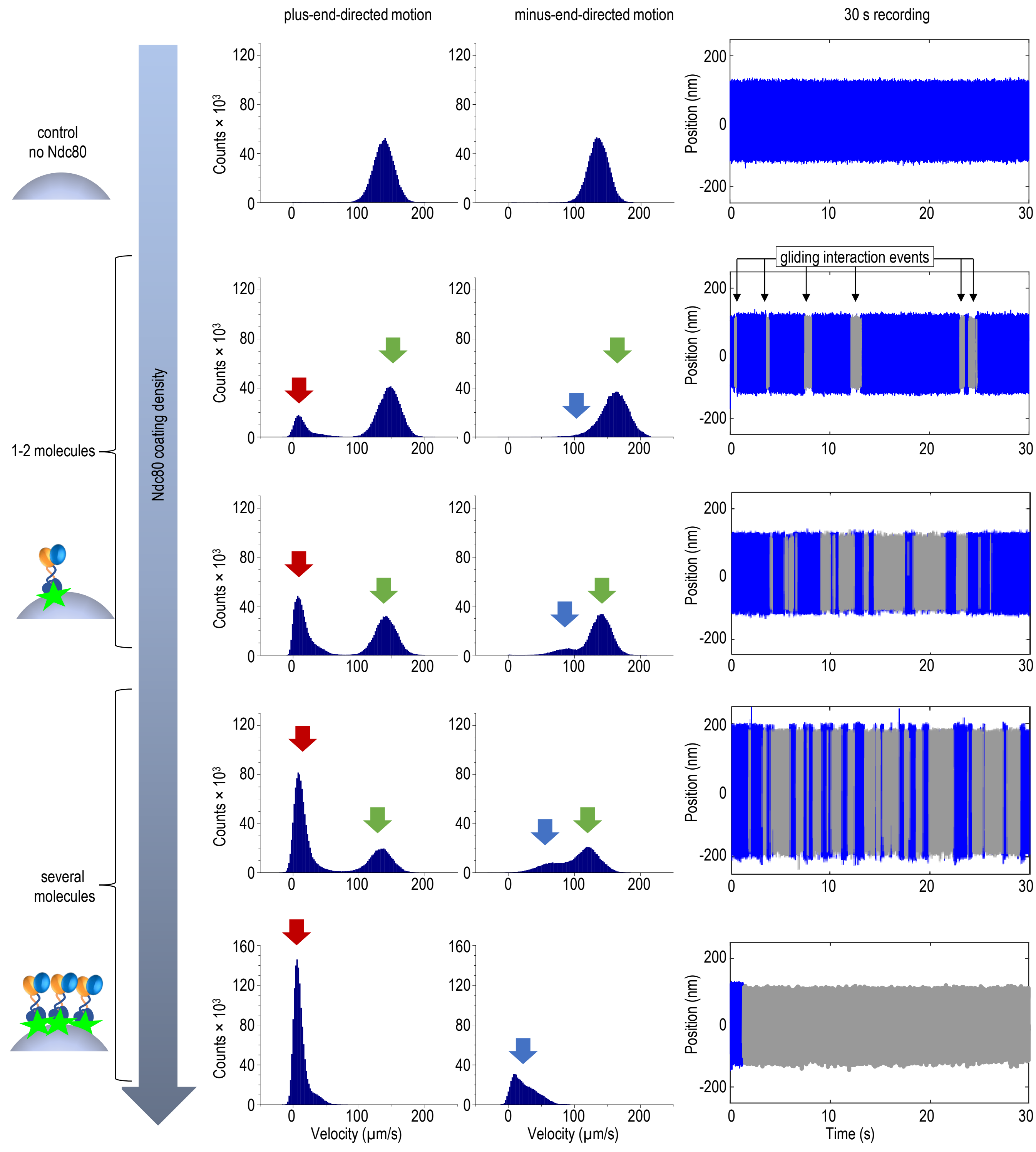


**Fig. S9. Example recordings of pedestals coated with Ndc80c at varying densities.** Each row shows histogram distributions of dumbbell bead velocities in two directions, based on 30 s position recordings at 4 pN force (right); Ndc80c sliding interaction events are shown in gray. Green arrows indicate free velocity peaks, red arrows mark velocity peaks for the plus-end-directed Ndc80c sliding, and blue arrows denote the approximate position of minus-end-directed sliding peaks. At low density of Ndc80c (< 20% interacting pedestals), the free velocity and plus-end sliding peaks are distinct, while the minus-end direction typically exhibits a single, slightly skewed peak. As Ndc80c density increases, the plus-end free velocity peak disappears, indicating near-continuous sliding, while the minus-end-directed free velocity peak is replaced by a peak with visibly lower velocity.

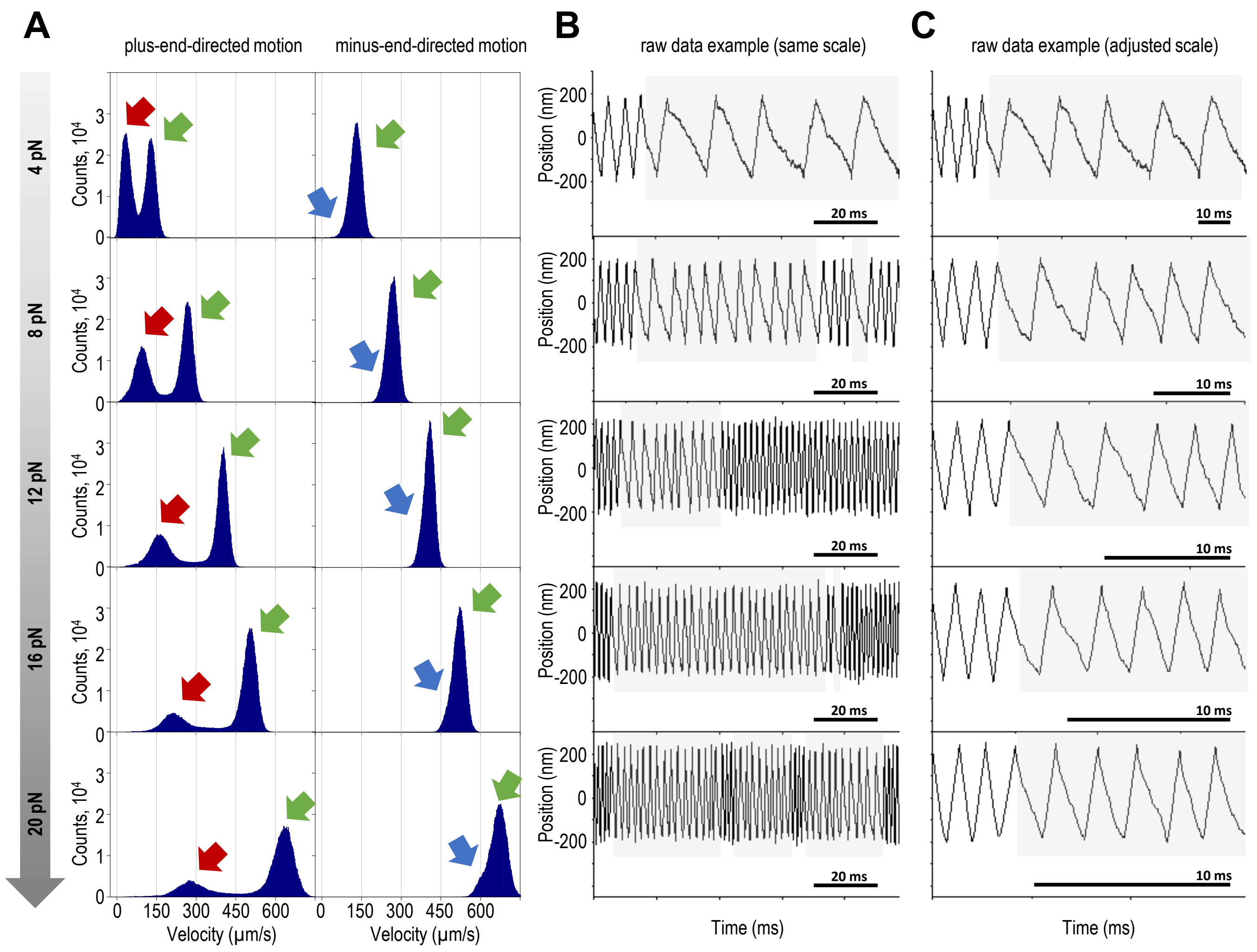


**Fig. S10. Single Ndc80c molecules exhibit asymmetric sliding across a range of dragging forces. (A)** Histogram distributions of instantaneous velocities for pedestals coated with Ndc80c Bonsai at varying forces. As the dragging force increases, velocity peaks shift rightward, indicating faster sliding. Distinct velocity peaks are not visible for minus-end-directed sliding due to its higher speed. (**B**) Example recordings of dumbbell bead positions shown on the same scale. (**C**) Magnified fragments of the recordings from panel B, presented with an adjusted scale to enable visual comparison.

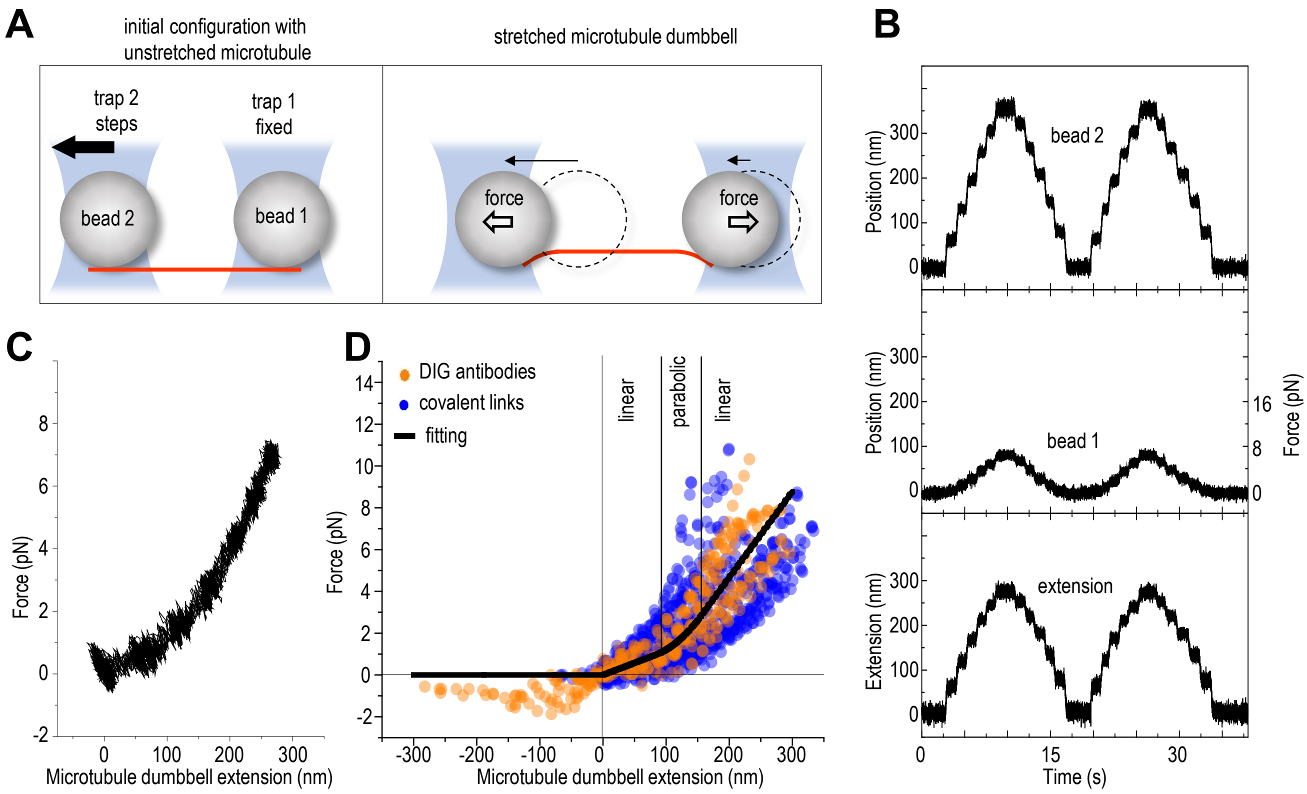


**Fig. S11. Stretching of the microtubule dumbbells. (A)** A schematics of the experiment to determine microtubule dumbbell compliance, see Supplementary text, Theoretical modelling Part 2 "Theoretical description of the ultrafast force-clamp assay". Microtubule dumbbell was stretched by moving trap 2 in 75 nm steps, while trap 1 was stationary. Thin arrows denote bead displacements from initial positions (dashed circles). **(B)** Example experimental data showing beads coordinates and measured extension, for traps stiffness 0.08 pN/nm, sampling rate 5 kHz. **(C)** Force-extension for experiment in panel B. **(D)** Force-extension data points for microtubule dumbbells that used DIG-labeled tubulin. Different colors show data for dumbbell beads that were coated with sheep anti-DIG antibodies using two different approaches: via bead-immobilized anti-sheep antibodies (N = 8 dumbbells) or via direct linkage to carboxylated beads (N = 11). Black curve is the experimental force-extension dependency which was constructed via piece-wise fitting of two linear and one parabolic regions, see Supplementary text.

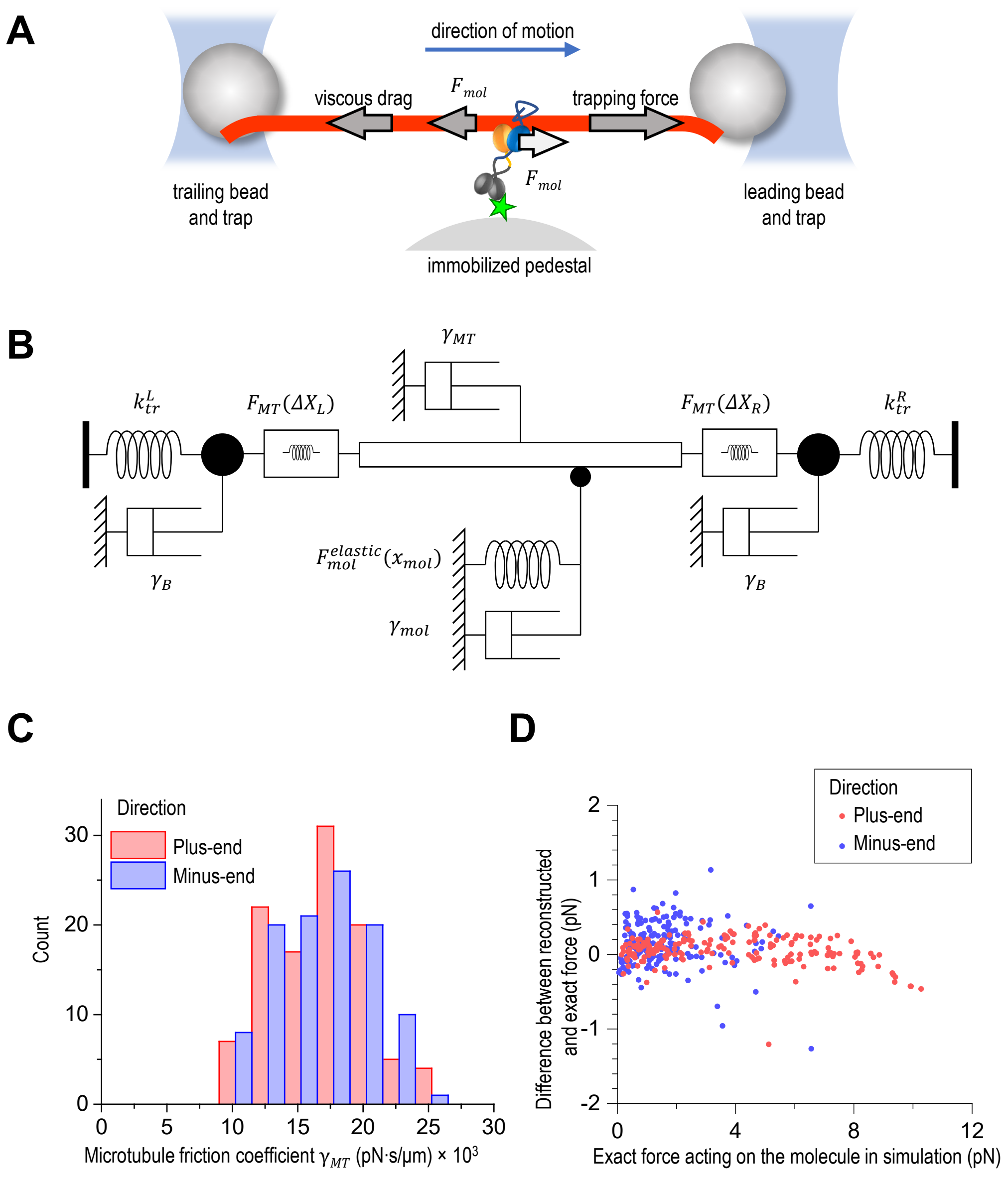


**Fig. S12. Theoretical modeling of the UFFC assay. (A)** Key elements of the UFFC experimental system. Gray arrows depict forces acting on the microtubule. **(B)** Mechanical one-dimensional representation of the UFFC experimental system. Dashpots symbolize viscous drag acting on all elements, traps are represented with Hookean springs, whereas the dumbbell and molecule compliance are modeled with nonlinear springs. (**C**) Estimation of microtubule friction coefficient. Two overlapping histograms display the friction coefficients for microtubules used in our UFFC experiments with Ndc80c Bonsai, bin width 0.001 pN·s/µm. (**D**) The force acting on the sliding Ndc80c molecule in the two-site model was determined using two methods. The "exact force," shown on the X-axis, was directly derived as the model output, with each point representing the result of a single simulation under a clamped force of 2–12 pN and parameter values specified in Tables S3 and S4. The Y-axis displays the difference between this exact force and the force reconstructed using our algorithm for analyzing experimental data (see eq. 16).

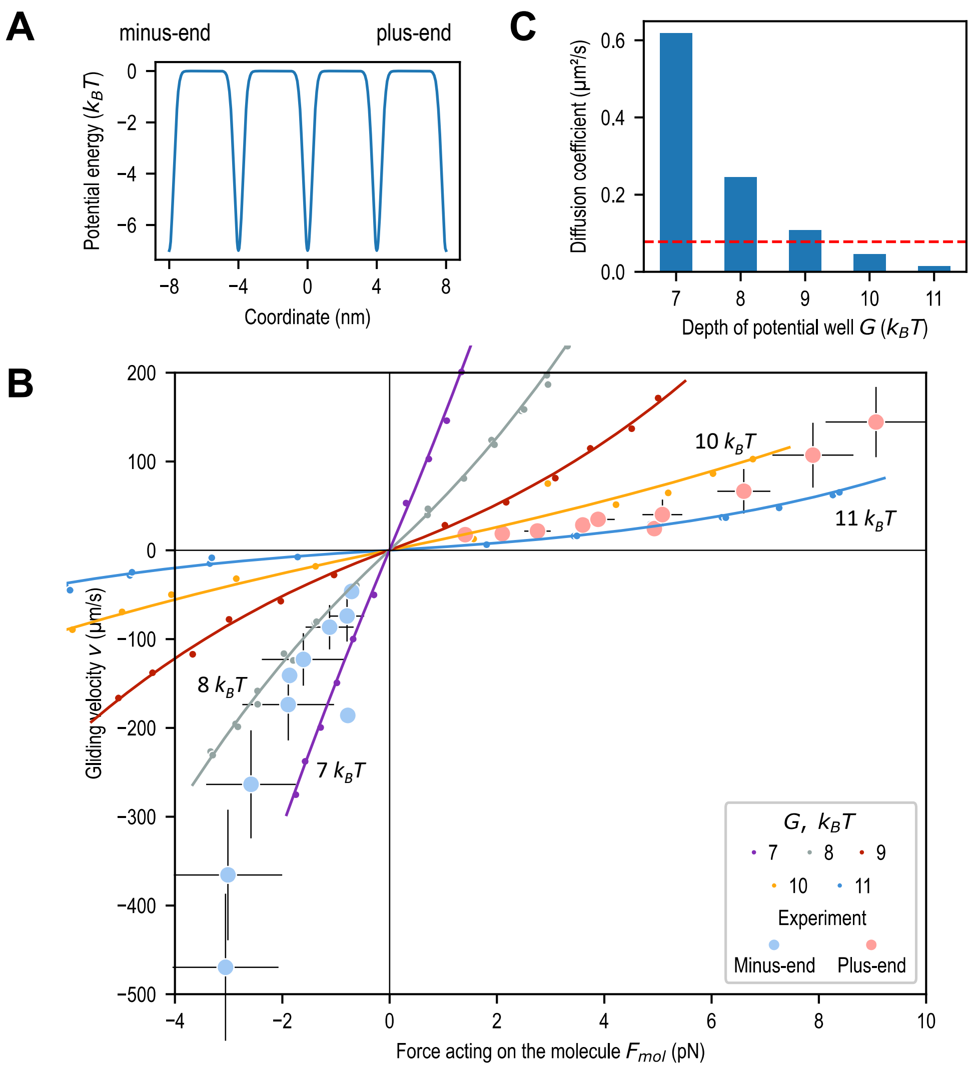


**Fig. S13. Single-site model for Ndc80c interaction with microtubule. (A)** Energy profile of a periodic potential landscape with a well depth of $G = 7\ k_BT$. **(B)** Predicted force-velocity dependencies. Each point represents one simulation, solid lines are exponential fittings. Large blue and pink dots correspond to binned experimental results, shown with SD. **(C)** Diffusion coefficients estimated from the initial slopes of the force-velocity dependencies in panel B. Red dashed line corresponds to the previously reported diffusion coefficient of Ndc80c: $D_{Ndc80} = 0.078\ \mu\text{m}^2/\text{s}$ (*33*).

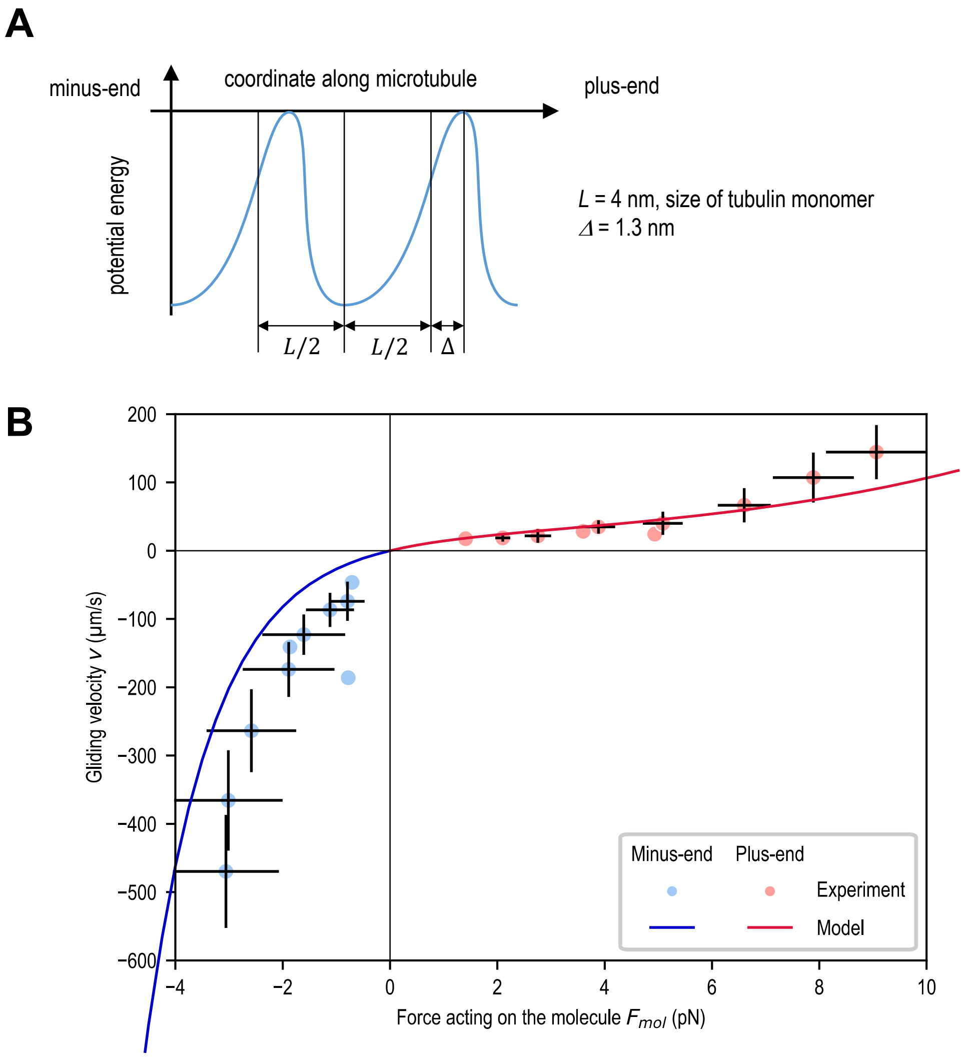


**Fig. S14. Asymmetric transition-state model for Ndc80c interaction with microtubule. (A)** Asymmetric energy potential for molecular translocation, here *L* denotes the period of potential well lattice and Δ denotes the asymmetry parameter. **(B)** The best-fit approximation of experimental velocities is with the asymmetry parameter Δ = −1.3 ± 0.1 nm (solid line).

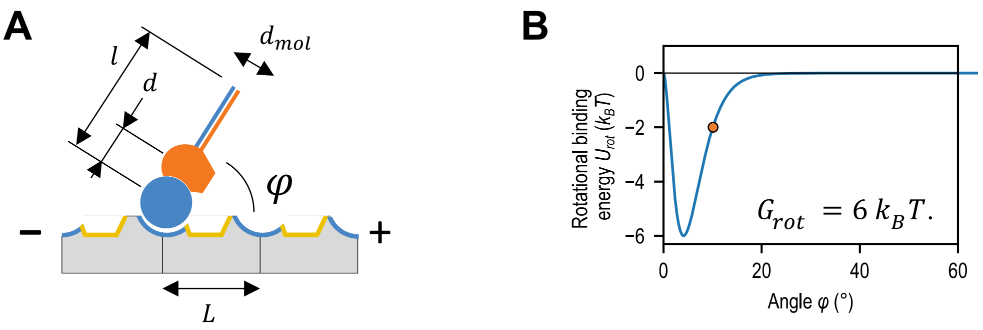


**Fig. S15. Two-site model for Ndc80c interaction with microtubule. (A)** The Ndc80c molecule is modeled as a rigid rod containing two microtubule-binding sites (corresponding to the Hec1 and Nuf2 CHDs). In the absence of force, the rod equilibrates at a $\varphi_0 = 60°$ angle relative to the microtubule axis. (**B**) Potential energy of interaction between the Nuf2 site and microtubule ($U_{rot}$) as a function of the rotation angle. For example, $U_{rot} = 2\ k_BT$ for $\varphi \approx 10°$ (depicted with an orange dot).

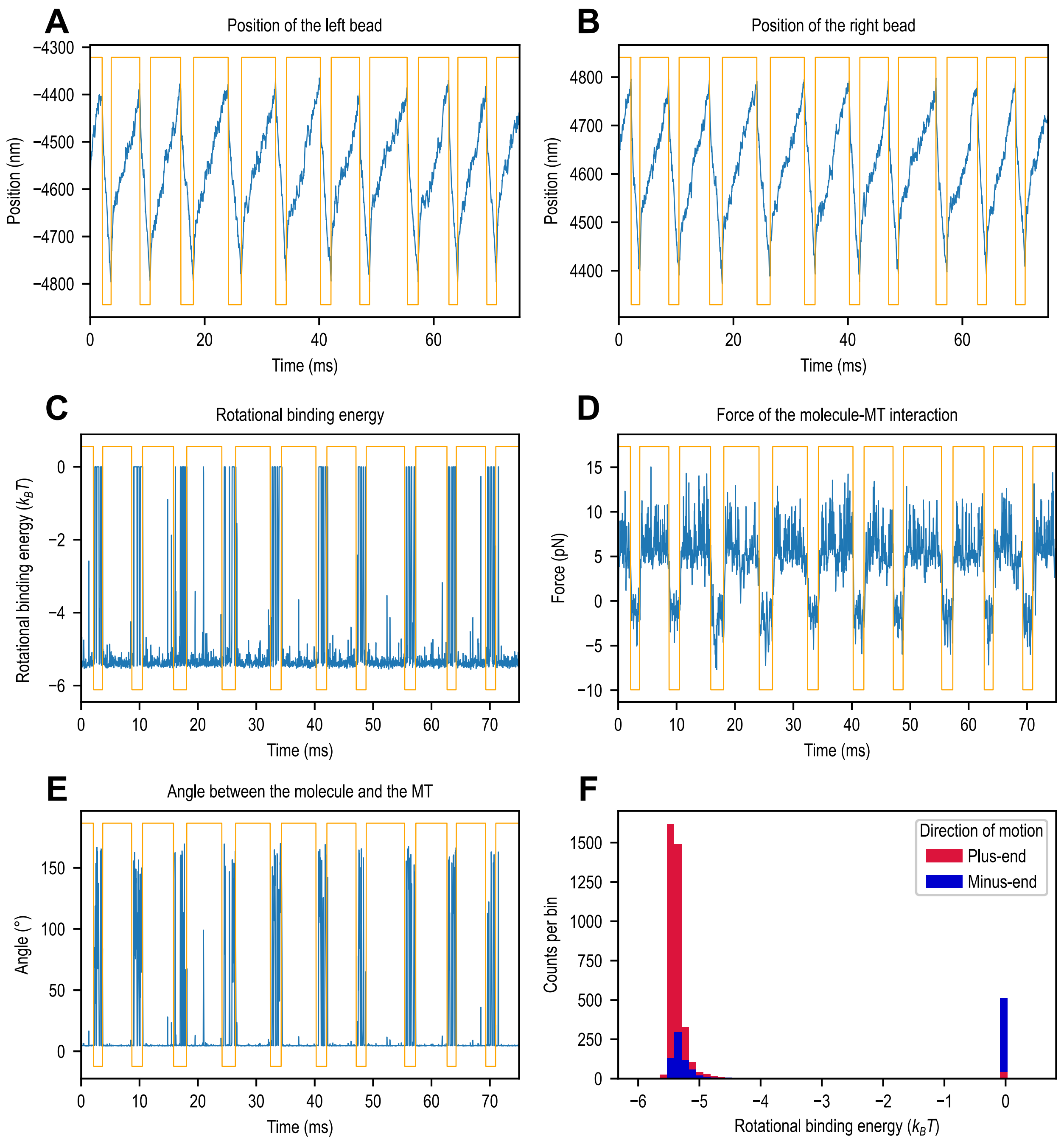


**Fig. S16. Typical output from a simulation using the two-site model.** Simulations were performed with the following parameters: $G_{main} = 6\ k_BT$, $G_{rot} = 6\ k_BT$, $k_{rot} = 10^{-2}$ pN·µm and a clamped force of $F = 8$ pN. Only 75 ms of the calculated results are displayed. Blue lines represent raw simulation data averaged over 15 µs, while orange lines depict directional changes in force. The top of each orange line indicates movement toward the microtubule's plus-end, while the bottom indicates motion toward the minus-end. **(A), (B)** Changes in positions of the left and right dumbbell beads, $x_B^L$ and $x_B^R$. **(C)** Rotational binding energy of the second site, $U_{rot}$. **(D)** Force acting on the Ndc80c molecule, $F_{mol}$. **(E)** Changes in the angle $\varphi$ between the Ndc80s rod and the microtubule. **(F)** Histogram of the second-site binding energy during the simulated segment (refer to panel C).

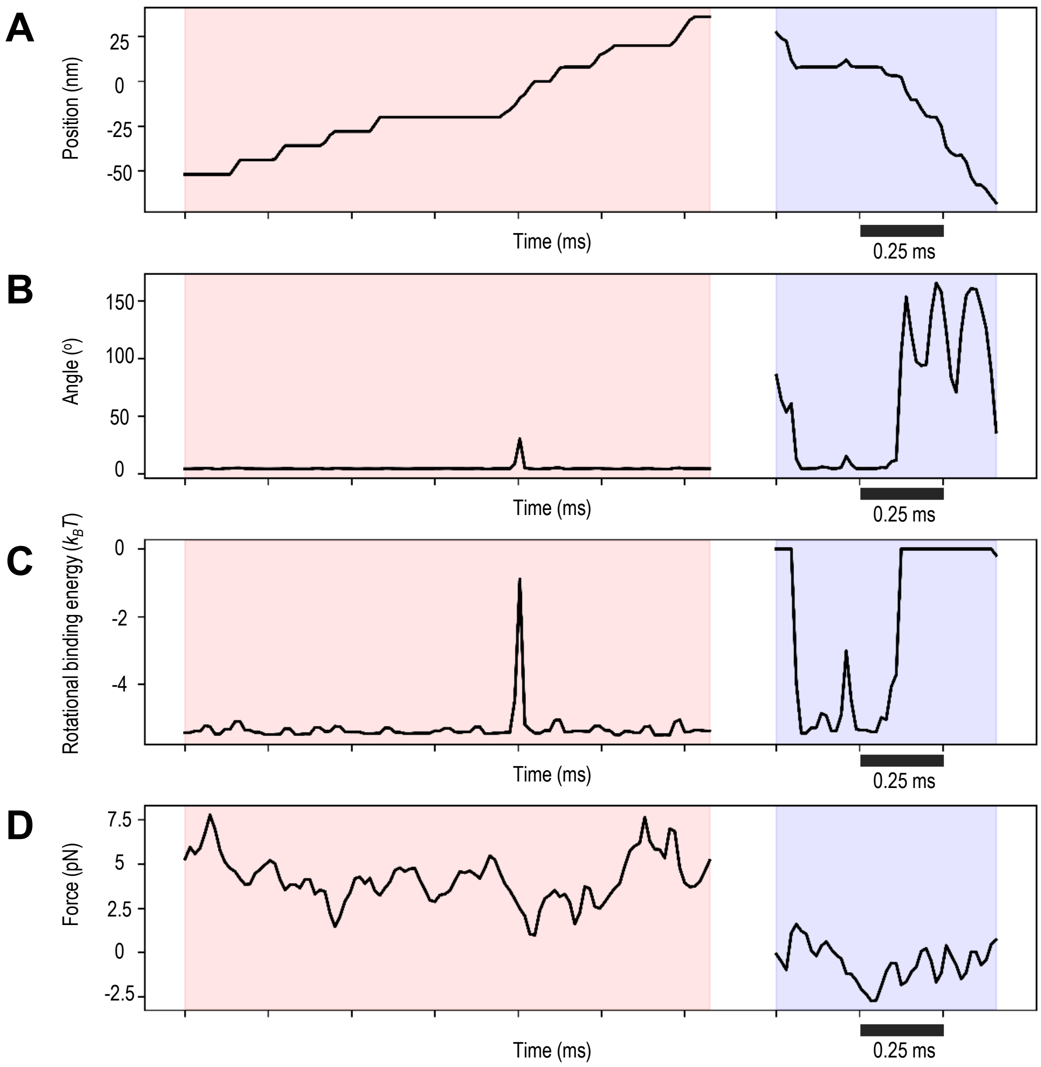


**Fig. S17. Numerical output from the simulation shown in Video 1.** Simulation results corresponding to Video 1. For detailed parameter and output descriptions, refer to the legend for fig. S16 and Video 1. Pink color – plus-end-directed sliding, blue color – minus-end-directed sliding. Although the simulation was performed at 4 pN force applied to the dumbbell, the average force experienced by the Ndc80c molecule is lower due to the viscous friction of the dumbbell. Notably, the force magnitude fluctuates significantly in either pulling direction, requiring continuous adaptation by the Ndc80c molecule.

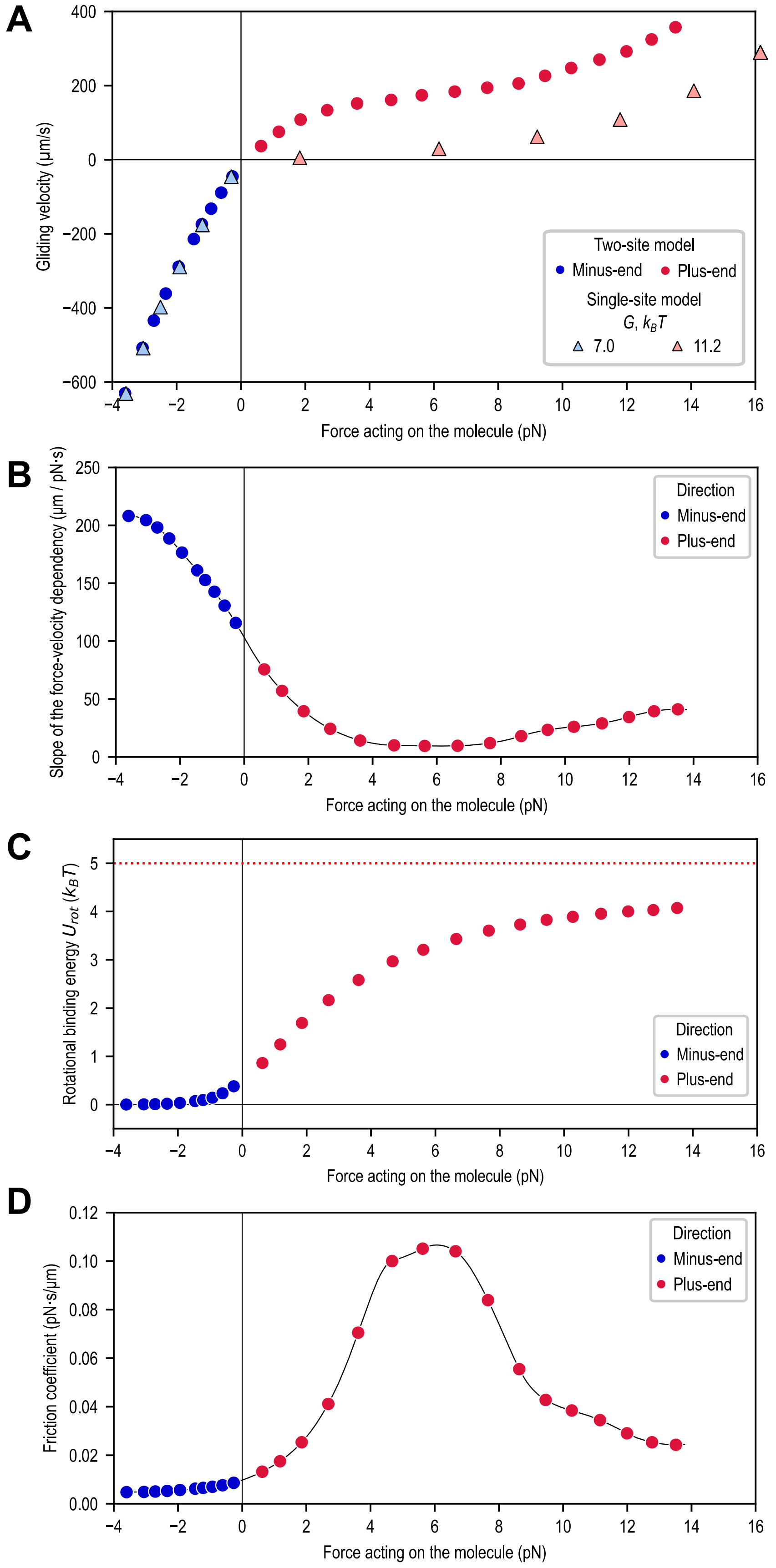


**Fig. S18. Analysis of the force-velocity relationship in the two-site model.** Simulations were performed using model parameters $G_{main} = 7\ k_BT$, $G_{rot} = 5\ k_BT$, $k_{rot} = 0.05$ pN · μm, $k_{tr}^{L,R} = 200$ pN/μm, and a simulation time of 150 ms. Red and blue dots represent results calculated with the same parameters but for opposite directions

**Fig. S18** (continued) of the clamped force. **(A)** Force velocity-dependency. Triangles indicate results from the single-site model. **(B)** Slope of the force-velocity dependency and a linear approximation near zero force (black line). **(C)** Rotational binding energy of the second site. **(D)** Friction coefficient, calculated as the reciprocal of the slope of the force-velocity dependency (panel B).

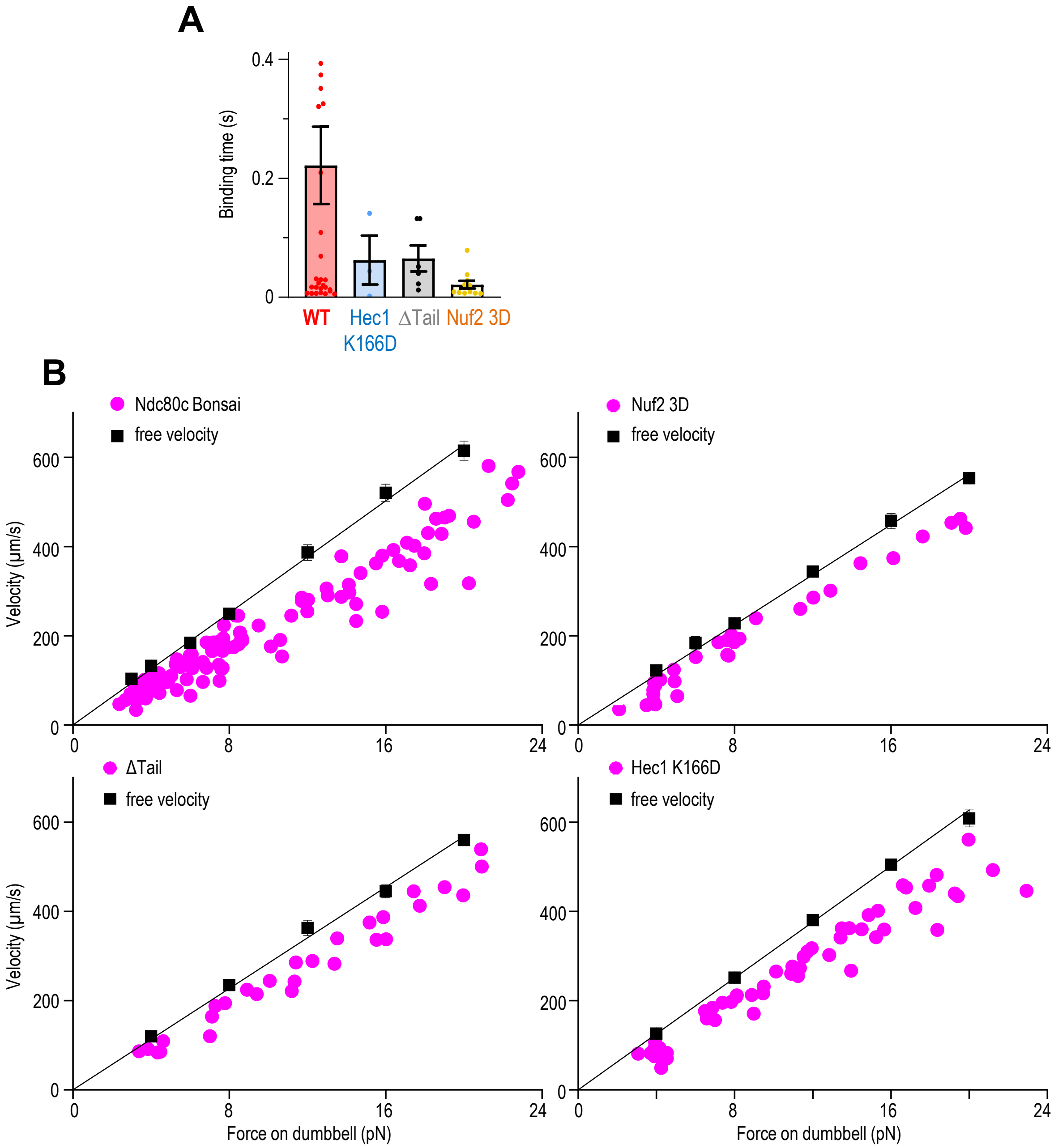


**Fig. S19. Experimental analysis of Ndc80c mutant proteins. (A)** Duration of sliding events. Dots show characteristic times for Ndc80c sliding in individual 30 s recordings under the 3-6 pN clamp force (see section 'Semi-automatic detection of continuous bi-directional Ndc80c sliding events'). Bars represent median values, whiskers represent 25%-75% interquartile range. Data points with time > 0.5 s are not shown (4 points for WT) for improved graph scaling. Duration of the sliding events is reduced for all Ndc80c mutant proteins. **(B)** The relationship between sliding velocity and force acting on a dumbbell is shown for various pedestal-immobilized proteins. Magenta dots represent the mean velocities of all minus-end-directed sweeps during interaction events in a single 30-second recording (with $n$ denoting the number of recordings and $N$ the number of experimental chambers): WT ($N$ = 43, $n$ = 106), 3D Nuf2 ($N$ = 12, $n$ = 30), Δ Tail ($N$ = 6, $n$ = 28), and Hec1 K166D ($N$ = 8, $n$ = 51). Black dots indicate the free velocity with SEM for the same experiments, while black lines are linear fits with a fixed zero intercept. For all mutants, the minus-end-directed velocity during interaction events was slower than the free dumbbell velocity, indicating friction generation. However, the similarity between the sliding velocities of the proteins and the free dumbbell velocity introduces substantial error when calculating molecular force, limiting the quantitative analysis of differences in the minus-end-directed velocities of these Ndc80 protein variants.
